# GlyConnect: a glycan-based conjugation extension of the GlycoDelete technology

**DOI:** 10.1101/2021.06.02.446789

**Authors:** Wander Van Breedam, Karel Thooft, Francis Santens, Sandrine Vanmarcke, Elise Wyseure, Bram Laukens, Berre Van Moer, Wim Nerinckx, Simon Devos, Annemieke Madder, Nico Callewaert

**Author notes:** These authors contributed equally (Wander Van Breedam, Karel Thooft).

## Abstract

Recently, our lab developed GlycoDelete, a technology suite that allows a radical simplification of eukaryotic N-glycosylation. The technology allows to produce glycoproteins that carry single GlcNAc, LacNAc, or LacNAc-Sia type glycans on their N-linked glycosylation sequons. GlycoDelete-type N-glycans are uniquely suited for glycan-based conjugation purposes, as these provide a short, homogeneous and hydrophilic link to the protein backbone. Targeting GlycoDelete-glycans allows for highly site-specific conjugation at sites in the protein which are normally occupied by bulky glycans, thus ensuring minimal interference with protein structure and function. The current manuscript describes the evaluation and optimization of both chemical and chemo-enzymatic conjugation of molecules onto the GlycoDelete-type glycans of a limited set of benchmark proteins.

## BACKGROUND

N-linked glycosylation is an important post-translational modification found on a whole range of proteins of eukaryotic origin, and occurs on asparagine residues in the sequence N-x-T/S (asparagine – any AA residue except proline – threonine or serine).^1–3^ N-glycosylation is initiated in the ER, and starts with a co-translational en-bloc transfer of a precursor glycan (GlcNAc_2_Man_9_Glc_3_) to the amide nitrogen of Asn.^2,4–6^ It has a strong link with the ER quality control system that prevents non-native proteins from being exported from the ER.^6^ This protein folding quality control happens through a folding-unfolding-refolding cycle (through the action of molecular chaperones, glycosidases and glycosyltransferases) that continues until a *de novo* synthesized protein adopts its native conformation.^1,2,4,5^ Once this box is ticked, the folded protein can continue its way through the secretory pathway. While the glycan heterogeneity in the ER is very limited, the concerted action of different glycosyltransferases, glycosidases, and other enzymes in the Golgi leads to further diversification or ‘maturation’ of the N-glycans.^2,4,5^ The variability inherent to this enzymatic process is at the basis of the highly diverse and complex N-glycan repertoires found on eukaryotic glycoproteins.

N-glycans play important structural and non-structural roles and are implicated in protein folding and solubility, protease resistance, masking of highly immunogenic protein stretches (‘glycan shielding’), and specific interactions with glycan-binding proteins.^5–7^ However, interestingly enough, many biopharmaceuticals don’t really need the N-glycans beyond protein folding.^8–11^ Inspired by this observation, we previously developed the GlycoDelete technology suite, wherein eukaryotic cells are hard-wired to yield glycoproteins decorated with small N-linked glycan stumps that are reminiscent of the non-reducing termini of (multi-branched) complex N-linked glycans found on mammalian glycoproteins.^4^ In its essence, GlycoDelete engineering is based on the heterologous expression of a Golgi-active endoglycosidase. Importantly, the ER-localized glycosylation household is left untouched, so as not to interfere with the protein folding and quality control processes.^4^ The engineering results in a drastic reduction in glycan heterogeneity, with the final N-glycan repertoire depending on the engineered host cell. GlycoDelete engineering of HEK293S cells resulted in an N-glycan repertoire mainly consisting of single N-acetylglucosamine (GlcNAc), N-acetyllactosamine (LacNAc), or sialylated LacNAc (LacNAc-Sia).^4^ GlycoDelete engineering of *Pichia pastoris (Komagatiella phaffii)* yielded a single GlcNAc N-glycan phenotype (unpublished results).

The GlycoDelete N-glycan stumps constitute an attractive handle for the bio-orthogonal site-specific conjugation of prosthetic groups like dyes, chelators, polymers, toxic payloads, etc. An advantage of this conjugation strategy is that the prosthetic groups end up in a space that is normally occupied by bulky N-glycans in the native glycoprotein, thus lowering the chance of interfering with protein structure and function. In this study, we evaluated several glycan-based conjugation strategies for GlycoDelete type glycans (GlyConnect). Both chemical and chemo-enzymatic conjugation approaches were optimized, that lead to the formation of a reactive aldehyde on the GlycoDelete glycan. The aldehyde is ultimately used for the conjugation of prosthetic groups on a small set of benchmark proteins, including a VHH (GFP-binding protein (GBP)-R86N), an antibody (obinutuzumab) and a growth factor (human granulocyte-macrophage colony-stimulating factor (hGM-CSF). Of note is that the focus lies on conjugation under mild conditions, i.e. aqueous medium and near-neutral pH. In parallel, the mammalian and yeast GlycoDelete strains were further engineered to increase compatibility with the GlyConnect methodology. The GlycoDelete-GlyConnect tandem presented here expands the range of conjugation strategies available for biopharmaceuticals and constitutes an attractive alternative to classical conjugation strategies.

## MATERIALS AND METHODS

### Reagents

#### Chemicals

Sodium periodate (NaIO_4_), p-phenylenediamine (PPD, 98% pure, solid), morpholine borane (MB, 95%) and aniline (99.5%, liquid), were purchased from Sigma-Aldrich. The purity of the NaIO_4_ stock was determined via standard KI/Na_2_S_2_O_3_ iodometric titration with starch as indicator^12^, and was determined to be 90%. UDP-GalNAz and Click-IT™ biotin DIBO alkyne were obtained from the GlyCLICK antibody labeling kit (Genovis). CMP-azido-sialic acid was purchased from R&D Systems (Bio-techne) and dibenzylcyclooctyne(DBCO)-PEG4-biotin from Jena Bioscience. Tris(2-carboxyethyl)phosphine (TCEP) was purchased from Gentaur.

#### Conjugates

Biotin-PEG_3_-ONH_2_ (EZ-Link™ alkoxyamine-PEG4-biotin) was obtained from Thermo Fisher Scientific, DTPA-NH_2_ (p-NH₂-Bn-CHX-A”-DTPA.2HCl.2H_2_O) was obtained from Macrocyclics, 5-kDa and 10-kDa linear PEG polymers (MeO-(CH_2_CH_2_O)_n_-NH_2_) were obtained from JenKem Technology (Custom Synthesis), 20-kDa PEG linear (CH_3_O-(CH_2_CH_2_O)n-CONH-CH_2_-O-NH_2_) and Y-shaped (CH_2_X-CHX-CH_2_-OOCNH-CH_2_ONH_2_ with X = (OCH_2_CH_2_)n-OCH_3_) were obtained from NOF Europe.

#### Enzymes

The galactose oxidase (GalOx) enzyme used in this study was produced in-house (Supplementary Information). The GalOx-F2 enzyme^13^ was provided by Prozomix as a cell-free extract containing 5% enzyme. Bovine liver catalase and horseradish peroxidase were obtained from Sigma Aldrich. Recombinant β-1,4-galactosyltransferase_Y289L (GalT-Y289L) was obtained from the GlyCLICK antibody labeling kit (Genovis)^8^, and recombinant human ST6Gal sialyltransferase 1/ST6GAL1 (aa 44-406) from R&D Systems (Bio-techne). FabRICATOR enzyme was purchased from Genovis.

### Production of benchmark GlycoDelete proteins

All benchmark proteins were produced in-house. GBP-R86N was produced in both HEK293- and Pichia-GlycoDelete strains. hGM-CSF and obinutuzumab were produced in the HEK293-GlycoDelete strain. Details about protein production and purification are provided as supplementary information.

### Exploring the glycan-based conjugation landscape

#### NaIO_4_-based conjugation on LacNAc-Sia-carrying glycoprotein

GBP-R86N protein (50 µg in 100µl; HEK293-GlycoDelete-produced) was oxidized with 10 mM NaIO_4_ in PBS buffer (pH 7) for 30 min on ice in the dark.^14–21^ Subsequently, excess NaIO_4_ was quenched by adding a 50-fold molar excess (vs NaIO_4_) of glycerol and incubating for 15 min on ice in the dark. Excess NaIO_4_, glycerol, and formaldehyde by-products were removed using PD-10 desalting columns (gravity flow protocol; PBS as equilibration buffer), and oxidized GBP-R86N protein was concentrated again using a Vivaspin 2 filter (3 kDa MWCO). Finally, oxime ligation was performed by adding a 100-fold molar excess of biotin-PEG_3_-ONH_2_ label (vs GBP-R86N protein) and slowly freezing the reaction mixture at −20°C.^22^

#### GalOx-based conjugation on LacNAc-carrying glycoprotein

A mixture of 200 µl GBP-R86N protein (Pichia-GlycoDelete-LacNAc-produced; 7.4E-8 moles), 18 µl horseradish peroxidase (final concentration 5U/ml), 45 µl bovine liver catalase (final concentration 200 U/ml), 58 µl GalOx (58 U; 7.4E-10 moles), 6.5 µl biotin-PEG_3_-ONH_2_ label (7.4E-6 moles), 200 µl 0.5 M NaP_i_ buffer (pH 7) and 472.5 µl H_2_O was incubated for 19h in the dark at 20 °C, followed by 44h in the dark at 4°C.^23^

#### Click-based conjugation on LacNAc-carrying glycoprotein

GBP-R86N glycoprotein was modified with azido-sialic acid (AzSia), by combining 10 µl GBP-R86N protein (Pichia-GlycoDelete-LacNAc-produced; 5 µg/µl), 3.64 µl ST6GAL1 protein (0.549 µg/µl), 7.41 µl CMP-azido-sialic acid stock (1 mM), 1.09 µl NaCl stock (5 M) and 27.86 µl of 25 mM TBS buffer (pH 7.4). The reaction was incubated for 1h at 37°C and 12h at 30°C in the dark. To react the azide function with DBCO-PEG4-Biotin, 1.76 µl DBCO-PEG4-biotin label (26.67 mM stock) was added to a 30 µl sample of the AzSia-modified GBP-R86N. The reaction mixture was incubated for 17h at 20 °C in the dark.

#### Click-based conjugation on GlcNAc-carrying glycoprotein

The protocol of the GlyCLICK antibody labeling kit was adapted as followed. GBP-R86N glycoprotein was modified with azido-galactose (GalNAz) by combining 5 µl GBP-R86N protein (Pichia-GlycoDelete-produced; 5 µg/µl), 12.5 µl GalT-Y289L, 15 µl of buffer additive and 167.5 µl of 25 mM TBS buffer (pH 7.4). The mixture was added to a vial containing 0.11 mg UDP-GalNAz and incubated for 20h at 30°C in the dark. Excess UDP-GalNAz was removed using a Vivaspin 2 filter (3 kDa MWCO) and the buffer exchanged to 25 mM TBS buffer (pH 7.4). The final volume was corrected to 250 µl. To react the azide function with biotin-DIBO label, the Click-IT™ biotin DIBO alkyne label provided in the GlyCLICK antibody labeling kit was reconstituted in 25 µl DMSO, and 12.5 µl label was added to the GalNAz-modified GBP-R86N. The reaction mixture was incubated for 18h at 25°C in the dark.

#### GalOx-F2-based conjugation on GlcNAc-carrying glycoprotein

A mixture of 200 µl GBP-R86N protein (Pichia-GlycoDelete-produced; 7.4E-8 moles), 18 µl horseradish peroxidase (final concentration 5 U/ml), 45 µl bovine liver catalase (final concentration 200 U/ml), 21 µl GalOx-F2 (7.4E-10 moles), 6.5 µl biotin-PEG_3_-ONH_2_ label (7.4E-6 moles), 5 µl CuSO_4_.5H_2_O (final concentration 0.05 mM), 200 µl 0.5 M NaP_i_ buffer (pH 7) and 504.5 µl H_2_O was incubated for 19h in the dark at 20 °C, followed by 44h in the dark at 4°C.^13^ As a control, an identical GalOx-F2-based reaction was performed on Pichia-GlycoDelete-LacNAc-produced GBP-R86N protein.

### Optimization of selected conjugation strategies

#### Optimization of NaIO_4_-based oxidation

Sialic acid-specific oxidation with NaIO_4_ was optimized on a HEK293-GlycoDelete-produced GBP-R86N protein stock, which mainly contained non-glycosylated and LacNAc-Sia glycosylated species. The non-sialylated GBP-R86N species in the mixture served as internal control for undesired side-oxidation/conjugation. Briefly, GBP-R86N protein was oxidized with varying concentrations of NaIO_4_ (0.5, 1, 2.5, 5 and 10 mM) in a 0.1 M NaP_i_ buffer (pH 7) on ice and in the dark for 30 min. Subsequently, the excess NaIO_4_ was quenched with 50 eq. of glycerol (vs NaIO_4_) and reactions were incubated for 15 min on ice and in the dark. Then excess NaIO_4_, glycerol and formaldehyde by-product were removed using Pierce™ Polyacrylamide Spin Desalting Columns (7K MWCO, 0.7 mL) and the buffer exchanged to 0.1 M NaP_i_ buffer (pH 7). Eventually, a reaction of 30 min at 4°C in the dark using 10 mM NaIO_4_ in 0.1 M NaP_i_ (pH 7) was selected as the optimized condition for LacNAc-Sia-specific NaIO_4_-based glycoprotein oxidation.

#### Optimization of GalOx-based oxidation

Galactose-specific oxidation with GalOx was optimized on a Pichia-GlycoDelete-LacNAc-produced GBP-R86N protein stock, which mainly contained non-glycosylated, GlcNAc-glycosylated and LacNAc-glycosylated species. The non-galactosylated GBP-R86N species in the mixture served as internal control for undesired side-oxidation/conjugation. Considering that GalOx uses O_2_ as a substrate, reactions were performed in closed vessels with ample headspace to include sufficient oxygen. To determine the optimal reaction temperature, 20 µg GBP-R86N was combined with a fixed amount of GalOx enzyme (0.01 molar equivalent; 2.3U) and 50eq. of biotin-PEG_3_-ONH_2_ label in a 0.1 M NaP_i_ buffer (pH 7), and incubated for 16h at 4°C or 20°C. To determine an optimal GalOx concentration, 20 µg GBP-R86N was mixed with a 25-fold molar excess of biotin-PEG_3_-ONH_2_ label and different amounts of GalOx (0, 1, 2, 4, or 8U) in a 0.1 M NaP_i_ buffer at pH 7. The reaction mixtures were incubated for 5h at 20 °C, upon which para-phenylene diamine (PPD) was added to a final concentration of 1 mM and reactions were further incubated for 2h at 20 °C. A reaction of 5h at 20°C using 2U of GalOx for 1.6 nmol of protein (ca. 0.01 eq.) in a 0.1 M NaP_i_ (pH 7) was selected as the optimized condition for LacNAc-specific GalOx-based glycoprotein oxidation.

#### Optimization of oxime ligation-based conjugation

Oxime conjugation was optimized on NaIO_4_-oxidized HEK293-GlycoDelete-produced GBP-R86N in a 0.1 M NaP_i_ buffer pH 7, using biotin-PEG_3_-ONH_2_ as the label. Temperature (4°C, 20°C, or −20°C (slow freezing)), label equivalents (25, 50 or 100 eq.), reaction time (30 min to 20h), salt concentration (0, 1, 2, or 3 M), and catalyst (different concentrations of aniline or PPD) were varied. A reaction of 2h at 20°C using 25 eq. label and 1 mM of PPD in a 0.1 M NaP_i_ buffer (pH 7) was selected as the optimized condition for oxime ligation-based glycoprotein conjugation.

#### Optimization of reductive amination-based conjugation

Reductive amination was optimized on NaIO_4_-oxidized HEK293-GlycoDelete-produced GBP-R86N, using aniline (50 eq.) as the label and morpholine borane (35 mM) as the reducing agent. Protein samples were buffer exchanged to either a 0.1 M NaP_i_ buffer at pH 7 or pH 6, or to 0.1 M NaAc buffer at pH 5 or pH 4, upon which label and reducing agent were added and reactions were incubated for 4h at 20 °C. A reaction of 4h at 20°C using 50 eq. label and 35 mM morpholine borane in a 0.1 M NaAc buffer (pH 5) was selected as the optimized condition for reductive amination-based glycoprotein conjugation.

### Expansion of the GlyConnect space

#### PEG polymer conjugation on GBP-R86N

Oxime-based conjugation of larger PEG polymers was performed on NaIO_4_-oxidized, HEK293-GlycoDelete-produced GBP-R86N. The non-sialylated GBP-R86N species in the mixture served as internal control for undesired side-oxidation/conjugation. For oxidation, the optimized condition for LacNAc-Sia-specific NaIO_4_-based glycoprotein oxidation was used. For conjugation, the optimized condition for oxime ligation-based glycoprotein conjugation was used, but different concentrations of the different PEG-aminooxy polymers (5 kDa and 10 kDa PEG; 25 or 10 eq.) and PPD (25, 5 or 1 mM) were tested. Additional conjugations were performed for linear and Y-shaped 20 kDa aminooxy PEG chains, using the optimized oxidation and conjugation conditions.

#### DTPA-NH_2_ conjugation on GBP-R86N

Reductive amination-based DTPA-NH_2_ conjugation was performed on NaIO_4_-oxidized, HEK293-GlycoDelete-produced GBP-R86N. The non-sialylated GBP-R86N species in the mixture served as internal control for undesired side-oxidation/conjugation. For oxidation, the optimized condition for LacNAc-Sia-specific NaIO_4_-based glycoprotein oxidation was used. For conjugation, the optimized condition for reductive amination-based glycoprotein conjugation was used, but the amount of label was increased to a 100-fold molar excess and reactions were incubated overnight. Control reactions with non-oxidized GBP-R86N or without the addition of label and reducing agent were included in the experimental setup.

#### PEG polymer conjugation on hGM-CSF

Oxime-based conjugation of larger PEG polymers was performed on NaIO_4_-oxidized, HEK293-GlycoDelete-produced hGM-CSF. For oxidation, the optimized condition for LacNAc-Sia-specific NaIO_4_-based glycoprotein oxidation was used. Control reactions without the addition of NaIO_4_ were included in the experimental setup. After buffer exchange to either a 0.1 M NaAc buffer (pH 5), a 0.1 M NaP_i_ buffer (pH 6) or a NaP_i_ buffer (pH 7) containing 1 mM PPD, an oxime ligation reaction was performed in the presence of 50 eq. of biotin-PEG_3_-ONH_2_ or 50 eq. of 10 kDa PEG-ONH_2_ polymer for 2h at 20 °C.

#### Oxime-based conjugation on obinutuzumab

Oxime-based conjugation was performed on HEK293-GlycoDelete-produced obinutuzumab. For oxidation, the optimized condition for LacNAc-Sia-specific NaIO_4_-based glycoprotein oxidation was used, but the oxidation buffer was changed to a 0.1 M NaP_i_ buffer (pH 6.5) and a range of NaIO_4_ concentrations (0.1, 1, 5 or 10 mM) was tested. After buffer exchange to a 0.1 M NaP_i_ buffer (pH 7), ligation was performed according to the optimized condition for oxime ligation-based conjugation.

### Protein analytics

#### LC-MS

Protein samples (10 µg; ca. 10-20 µl injection volume depending on protein concentration) were injected on a Poroshell 300SB-C8 column (5 µm, 300Å, 1×75mm IDxL; Agilent Technologies) at a flow rate of 100 µl/min using solvent A (0.1% formic acid and 0.05% trifluoroacetic acid in water) and solvent B (0.1% formic acid and 0.05% trifluoroacetic acid in acetonitrile) as mobile phases. After on-column desalting for 2.5 min at 10% solvent B, the proteins were separated using a 6.5 min gradient from 10% to 90% solvent B. Column temperature was maintained at 60°C. Eluting proteins were directly sprayed in the mass spectrometer with an ESI source using the following parameters: spray voltage of 4.2 kV, surface-induced dissociation of 30 V, capillary temperature of 325 °C, capillary voltage of 35 V and a sheath gas flow rate of 7 (arbitrary units). The mass spectrometer was operated in MS1 mode using the orbitrap analyzer at a resolution of 60,000 (at m/z 400) and a mass range of 600-4000 m/z, in profile mode. The resulting MS spectra were deconvoluted with the BioPharma Finder^TM^ 3.0 software (Thermo Fischer Scientific) using the Xtract deconvolution algorithm (isotopically resolved spectra). The deconvoluted spectra were manually annotated.

In standard LC-MS analysis, samples were injected without prior clean-up/purification. When PEG polymers were conjugated, the samples were purified using Ni-NTA Spin Columns (Qiagen; Loading buffer: 10 mM imidazole in PBS, pH 7.5-8, Washing Buffer: 20 mM imidazole in PBS, pH 7.5-8 and Elution Buffer: 250 mM imidazole in PBS, pH 7.5-8) before LC-MS analysis. Antibody samples were pre-treated with FabRICATOR enzyme at 37°C for 2h (10 µg of antibody was incubated with 10U of FabRICATOR) and then incubated with TCEP at 37°C for 1h (final TCEP concentration at 10 mM).

#### SDS-PAGE and Western Blotting

Protein/conjugate samples were mixed with Laemmli sample buffer after which SDS-PAGE was performed on pre-cast polyacrylamide gels (PAGE) from Bio-Rad (4–20% Mini-PROTEAN® TGX™ Precast Protein Gels, 15-well, 15 µl, 130V). Upon Western blotting, 6xHis-tag- and Fc-specific Dylight 800-labeled detection antibodies (6X His Tag Antibody Dylight™ 800 Conjugated (VWR) and Human IgG (H&L) Antibody Dylight™ 800 Conjugated (tebu-bio)) were used to probe the nitrocellulose membranes and visualization was performed using an LI-COR Odyssey 2.1 infrared imaging system.

### GlycoDelete strain engineering

To generate the HEK293-GlycoDelete-SiaHigh strain, the HEK293-GlycoDelete cell line was modified by introducing a sialuria mutation in the UDP-N-acetylglucosamine-2-epimerase/N-acetylmannosamine kinase (GNE) gene through CRISPR-Cas9 editing. ^24^ The HEK293-GlycoDelete-LacNAc strain was generated by establishing a GNE gene knockout using CRISPR-Cas9. The Pichia-GlycoDelete-LacNAc strain was generated by introducing the human UDP-Glc-4-epimerase (GalE) gene and the human beta-(1,4)-galactosyltransferase (GalT) gene in the Pichia-GlycoDelete strain using a linearized pGAPNORMnn2SpGal10hGalT-I GlycoSwitch vector.^25^ Details are provided as supplementary information.

## RESULTS

### Exploring the glycan-based conjugation landscape

In a first set of experiments, we aimed to explore which glycan-based conjugation strategies were suited for conjugation to GlycoDelete type glycans (Supplementary Figure 9). To ensure maximal compatibility of the methodology with potentially fragile glycoprotein therapeutics, we restricted the screen to conjugation technologies that can proceed under mild conditions (aqueous, near-neutral pH). A glyco-engineered GFP-specific VHH protein containing a single N-glycosylation site (GBP-R86N) was chosen as the test case protein, as its small size and high degree of homogeneity render it an ideal benchmark for conjugation and accurate analysis, especially when produced in a GlycoDelete background. The GlycoDelete platform gives access to glycoproteins with terminal Sia, Gal, or GlcNAc residues, each of which can serve as a hinge for glycan-based conjugation.^4^ The strategies that were explored can be broadly divided in aldehyde-based reactions and azide-based reactions.

For conjugation to terminal Sia, classic (mild) NaIO_4_-mediated oxidation was used^14–16,18,20,21^, to selectively oxidize the glycerol side chain of the sialic acid residue, generating a reactive aldehyde on the GBP-R86N glycoprotein.^19,20,26,27^ Upon quenching the excess NaIO_4_ and removing the byproducts, the aldehyde was reacted with a small aminooxy-functionalized PEG chain with a terminal biotin via oxime ligation (biotin-PEG_3_-ONH_2_, Supplementary Figure 1). Oxidation/ligation proceeded fairly efficiently under the conditions used, with minimal or no side oxidation (Figure 1A).

**Figure 1.**
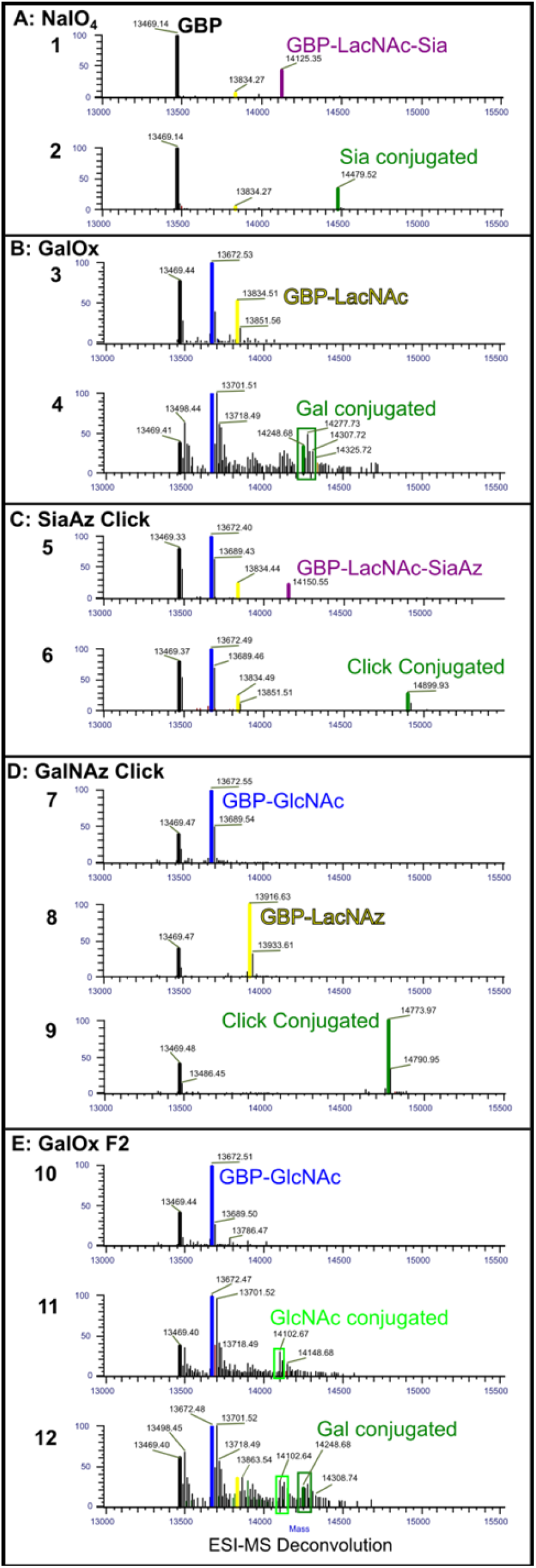
GBP-R86N conjugation using different glycan-specific conjugation methods. **A. NaIO_4_-based LacNAc-Sia conjugation**. HEK293-GlycoDelete-produced GBP-R86N starting material containing non-glycosylated (black, MW = 13469 Da), LacNAc (yellow, MW = 13834 Da) and LacNAc-Sia species (purple, MW = 14125 Da) (**A1**), was oxidized with NaIO_4_ followed by oxime conjugation of biotin-PEG_3_-ONH_2_ (**A2**). All of the sialylated fraction was conjugated successfully and selectively (green, MW = 14479 Da). **B. GalOx-based LacNAc conjugation.** Pichia-GlycoDelete produced GBP-R86N starting material containing non-glycosylated (black, MW = 13469 Da), GlcNAc (blue, MW = 13672 Da) and LacNAc species (yellow, MW = 13834 Da) (**B3**), was oxidized using GalOx, followed by oxime conjugation of biotin-PEG_3_-ONH_2_ (**B4**). Gal-selective conjugation (green, MW = 14248 Da) was achieved, but considerable side oxidation was observed (+16 Da). **C. Click-based LacNAc conjugation.** The LacNAc glycan of Pichia-GlycoDelete-produced GBP-R86N was enzymatically modified with an azido-sialic acid (**C5**) (purple, MW = 14150 Da) followed by click-reaction based conjugation of DBCO-PEG_4_-biotin (**C6**) (green, MW = 14899 Da). **D. Click-based GlcNAc conjugation.** The GlcNAc glycan of Pichia-GlycoDelete-produced GBP-R86N (**D7**), was enzymatically modified with an azido-galactose (**D8**) (yellow, MW = 13916 Da), followed by click-reaction based conjugation of Click-IT^TM^ Biotin DIBO Alkyne (**D9**) (green, MW = 14773). **E. GalOx-F2-based GlcNAc conjugation.** Pichia-GlycoDelete-produced GBP-R86N starting material containing non-glycosylated (black, MW = 13469 Da) and GlcNAc species (blue, MW = 13672 Da) (**E10**), was treated with GalOx-F2 enzyme followed by oxime conjugation of biotin-PEG_3_-ONH_2_ (**E11**). GlcNAc-selective conjugation was achieved but incomplete (light green, MW = 14086 Da) and side oxidation occurred (+16 Da). GalOx-F2 based oxidation and oxime conjugation in the presence of LacNAc glycan showed conjugation of both the GlcNAc and LacNAc glycan (**E12**). Both conjugations were incomplete and side oxidation occurred (+16 Da).

For conjugation to terminal Gal, recombinant GalOx was used to oxidize Gal at position C6, again generating an aldehyde and allowing specific conjugation with biotin-PEG_3_-ONH_2_ label.^27–29^ Oxidation/ligation proceeded efficiently, but significant side oxidation was observed (Figure 1B). In an alternative approach, we used recombinant human ST6Gal sialyltransferase 1 to derivatize the terminal Gal with an azido-sialic acid (AzSia) residue, which allows subsequent conjugation with alkyne-containing molecules in a click reaction.^30^ While the click reaction step was very efficient, the enzymatic addition of the AzSia residue was incomplete and showed room for further optimization (Figure 1C).

For conjugation to GlcNAc, recombinant GalT-Y289L was used to transfer an azide-modified Gal (GalNAz) to the GlcNAc residues, enabling conjugation through click chemistry.^31^ This approach proved to be highly specific and efficient (Figure 1D). In parallel, the use of a mutant GalOx (GalOx-F2)^13^ with a relaxed specificity was explored. This GalOx-F2 enzyme can oxidize GlcNAc at position C6, creating an aldehyde available for conjugation with the aforementioned aldehyde-reactive chemistries.^13^ This GalOx-F2-based approach on GlcNAc was clearly less efficient than the GalOx-based approach on Gal, as only a minor fraction of the GlcNAc glycan was conjugated with the aminooxy label (Figure 1E). This was not unexpected, as the GalOx-F2 mutant has been described to be much less efficient than the wild type GalOx enzyme.^13^

### Optimization of selected conjugation strategies

The Sia-specific conjugation strategy based on NaIO_4_ and the Gal-specific conjugation strategy based on GalOx were chosen for further optimization in the GlycoDelete context. In a first set of experiments, the oxidation reaction was optimized, i.e. the formation of the reactive aldehydes. In a second set of experiments, different aldehyde-based conjugation chemistries were evaluated and optimized.

### Oxidation reaction optimization

The standard protocol for Sia-specific oxidation of glycoproteins is the use of 1 mM NaIO_4_ in 0.1 M NaP_i_ buffer (pH 7) for 30 minutes and at 4°C in the dark.^14,16^ To start out, HEK293-GlycoDelete-produced GBP-R86N – a mixture of non-glycosylated GBP-R86N and GBP-R86N carrying a single LacNAc-Sia, LacNAc or GlcNAc type glycan – was subjected to this standard oxidation protocol, after which the excess NaIO_4_ was quenched and byproducts were removed. To ascertain formation of a reactive aldehyde, an excess amount of biotin-PEG_3_-ONH_2_ was added and the reaction mixtures were slowly frozen at −20°C to catalyze oxime formation.^22^ Surprisingly, LC-MS analysis revealed that the use of 1 mM NaIO_4_ resulted in only partial oxidation of the LacNAc-Sia. Further tests using a NaIO_4_ dilution series showed that increasing the NaIO_4_ content to 5 or 10 mM yielded complete oxidation of the LacNAc-Sia-carrying GBP-R86N. Importantly, these higher NaIO_4_ concentrations did not trigger side oxidation, as evidenced by the lack of reactivity of the non-sialylated species in the mixtures. Based on the data, the optimal condition for LacNAc-Sia-specific glycoprotein conjugation was 10 mM NaIO_4_ in 0.1 M NaP_i_ (pH 7) for 30 min at 4°C in the dark) (Figure 2A).

**Figure 2.**
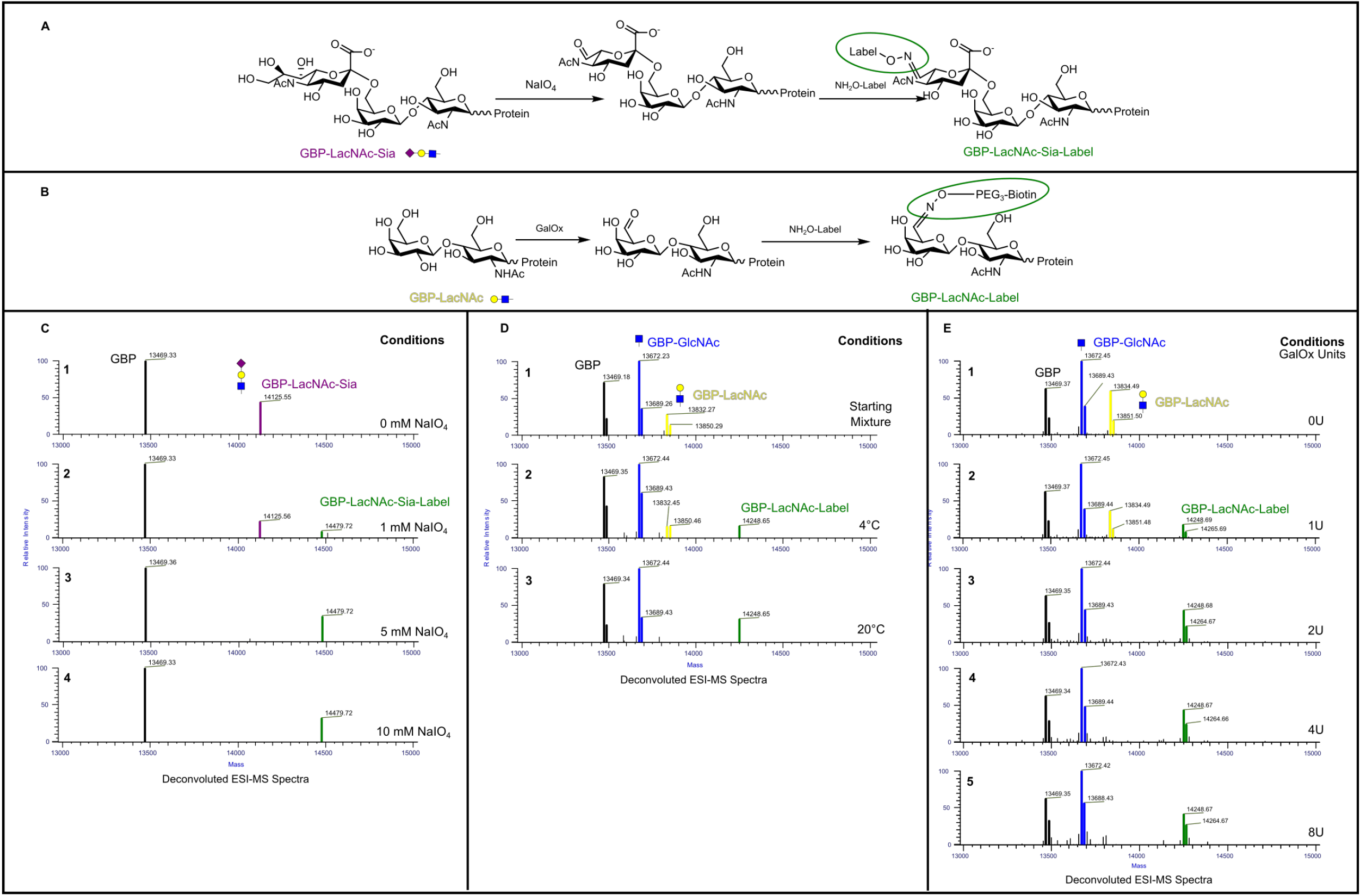
**A.** Reaction scheme of the LacNAc-Sia glycan oxidation using NaIO_4_, and subsequent oxime conjugation. **B**. Reaction scheme of the GalOx based oxidation of the LacNAc glycan and subsequent oxime conjugation. **C.** Deconvoluted MS spectra of GBP-R86N during NaIO_4_-based oxidation optimization. Oxidation was performed on HEK293-GlycoDelete-produced GBP-R86N, a mixture of non-glycosylated (black, MW = 13469 Da) and LacNAc-Sia species (purple, MW = 14125 Da) (**C1**). Conjugation of biotin-PEG_3_-ONH_2_ was performed at varying NaIO_4_ concentrations. With increasing NaIO_4_ concentrations, the LacNAc-Sia signal disappears and the conjugated species signal appears at the expected mass (green, MW = 14479 Da) (**C2**-**4**). **D.** Deconvoluted MS spectra of the GalOx-based oxidation of the LacNAc glycan on GBP-R86N and subsequent oxime conjugation of biotin-PEG_3_-ONH_2_, at different temperatures. Pichia-GlycoDelete-LacNAc-produced GBP-R86N, a mixture of non-glycosylated (black, MW = 13469 Da), GlcNAc (blue, MW = 13672 Da) and LacNAc species (yellow, MW = 13832 Da), was used (**D1**). The reaction was performed for 16h at 4°C (**D2**) and 20°C (**D3**). For each protein species in the mixture, a second MS signal was present (+17 Da), which corresponds to the non-pyroglutamate form. The conjugated species signal appears at the expected mass (green, MW = 14248 Da). **E.** Deconvoluted MS spectra of the GalOx-based oxidation of the LacNAc glycan on GBP-R86N and subsequent oxime conjugation of biotin-PEG_3_-ONH_2_, using different amounts of GalOx units. HEK293-GlycoDelete-LacNAc-produced GBP-R86N, a mixture of non-glycosylated (black, MW = 13469 Da), GlcNAc (blue, MW = 13672 Da) and LacNAc species (yellow, MW = 13834 Da), was used (**E1**). Again, for each protein species a second MS signal was present (+17 Da), which corresponds to the non-pyroglutamate form. The conjugated species signal appears at the expected mass (green, MW = 14248 Da) (**E2-4**).

For Gal-specific oxidation of glycoproteins, the use of GalOx was further explored. GalOx is an enzyme that catalyzes the oxidation of a range of primary alcohols, including D-Galactose, to the corresponding aldehyde with the reduction of molecular oxygen (O_2_) to hydrogen peroxide (H_2_O_2_).^23,32^ As is typical for enzymes, a direct relationship exists between temperature and GalOx activity. However, the fact that reactions are performed in aqueous medium and that GalOx uses O_2_ as a substrate complicates the matter, as there is an inverse relation between temperature and oxygen solubility in water.^32^ It was therefore imperative to evaluate the effects of the reaction temperature. In a first set of experiments, Pichia-GlycoDelete-LacNAc-produced GBP-R86N (20 µg) - a mixture of non-glycosylated GBP-R86N and GBP-R86N carrying a single LacNAc or GlcNAc type glycan - was combined with a fixed amount of GalOx enzyme (0.01 molar equivalent; 2.3U) and 50 eq. of biotin-PEG_3_-ONH_2_ (Supplementary Figure 1) and incubated for 16h at 4°C or 20°C. LC-MS analysis showed complete and specific conjugation – and thus complete oxidation – for the reaction performed at 20°C (Figure 2B). Although we observed incomplete conjugation for the reaction at 4°C, conjugation could be driven to completion by subsequent slow freezing, which strongly suggests that oxime formation kinetics were the limiting factor.^22^ Further experiments with a similar setup confirmed the data (Supplementary Figure 2).

To further standardize the GalOx-based oxidations, we performed an enzyme titration experiment. HEK293-GlycoDelete-LacNAc-produced GBP-R86N (20 µg) - a mixture of non-glycosylated GBP-R86N and GBP-R86N carrying a single LacNAc or GlcNAc type glycan - was mixed with a 25-fold molar excess of the biotin-PEG_3_-ONH_2_ label and different amounts of GalOx (0, 1, 2, 4, or 8U) in a 0.1 M NaP_i_ buffer (pH 7). The reaction mixtures were incubated for 5h at 20 °C, upon which PPD was added to a final concentration of 1 mM, and reactions were further incubated for 2h at 20 °C. The fact that PPD catalyzes oxime formation,^33^ but also blocks GalOx activity (own observation) makes it an ideal tool in these titration experiments. LC-MS analysis of the samples showed that 2U of GalOx was enough to achieve complete oxidation of the LacNAc-carrying GBP-R86N, without significant side oxidations (Figure 2C).

#### Conjugation reaction optimization

For the optimization of conjugation at (near-)neutral pH conditions, we used NaIO_4_-oxidized HEK293S-GlycoDelete-produced GBP-R86N as the substrate. The LacNAc-Sia-carrying fraction of the oxidized protein pool carries a reactive aldehyde, whereas the other glycoforms do not. Again, the non-sialylated proteins function as an internal control for conjugation specificity. Different aldehyde-reactive labels/tags/polymers were used (Supplementary Figure 1), in line with the specific conjugation chemistry envisioned.

To further explore oxime formation, we started out with the biotin-PEG_3_-ONH_2_ label we used before, as its small size (MW = 434 Da) and its chemical nature render it ideal for detection purposes and make it a good proxy for other small molecules of interest such as fluorescent tags or small cytotoxic drugs.

To evaluate the effects of the reaction temperature, NaIO_4_-oxidized GBP-R86N was mixed with a 100-fold molar excess of the biotin-PEG_3_-ONH_2_ label in a 0.1 M NaP_i_ buffer (pH 7) and incubated at 4°C, 20°C, or −20°C (slow freezing). LC-MS analysis showed that uncatalyzed oxime ligation occurs faster at 20°C than at 4°C. (Supplementary Figure 3) However, slow freezing at −20°C was by far the most efficient, with complete and selective conjugation being reached in a single freeze-thaw cycle (Figure 3).

**Figure 3.**
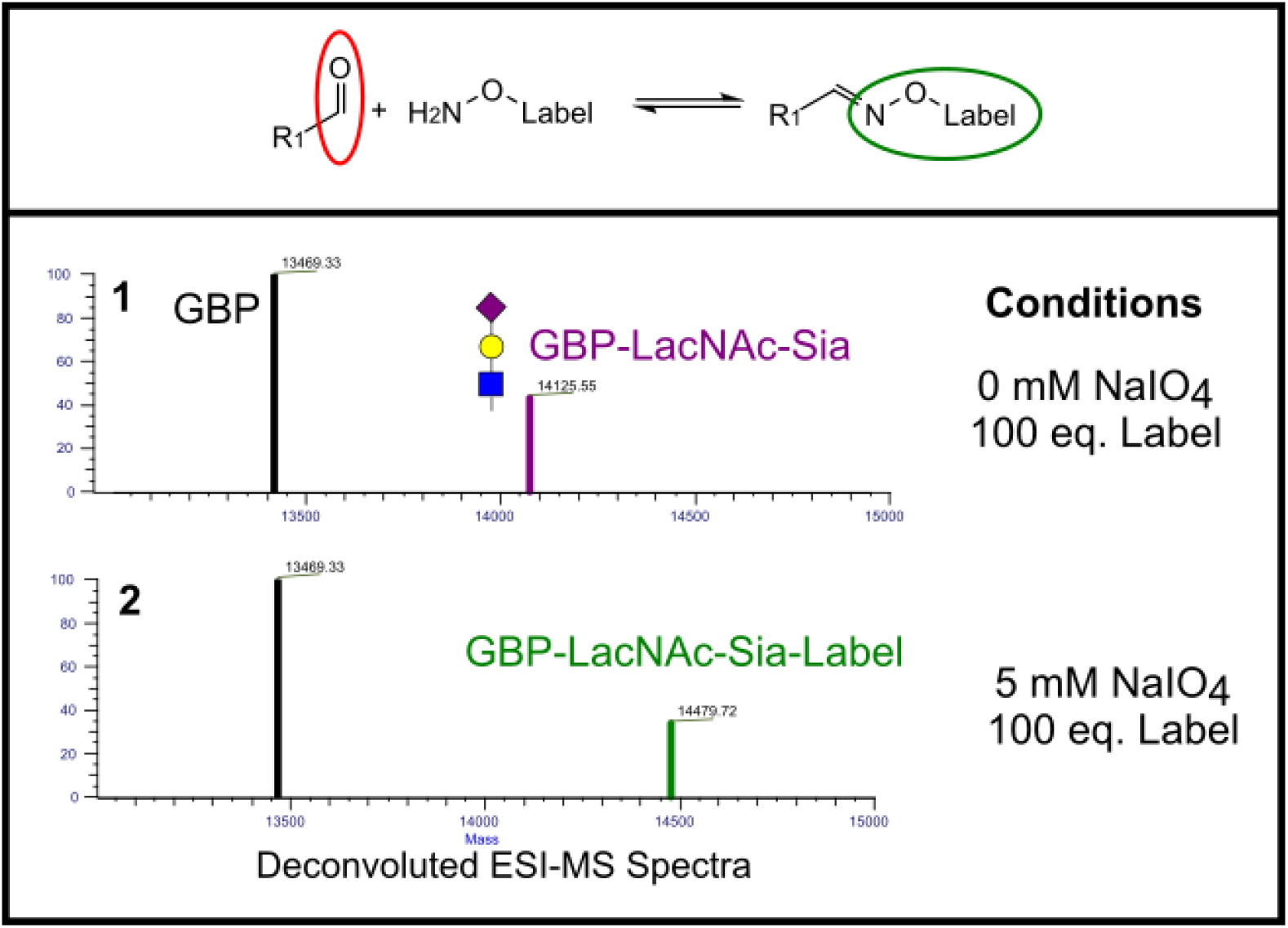
**Top**. Reaction scheme of oxime ligation. The aldehyde (red) originates from an oxidized glycan (R_1_) and reacts with an aminooxy-functionalized label to form the oxime conjugate (green). **Bottom.** Deconvoluted MS spectra of oxime conjugation on NaIO_4_-oxidized LacNAc-Sia glycans of GBP-R86N. The reaction was performed on HEK293-GlycoDelete-produced GBP-R86N, a mixture of non-glycosylated (black, MW = 13469 Da) and LacNAc-Sia species (purple, MW = 14125 Da) (1). The conjugated species signal appears at the expected mass (green, MW = 14479 Da) (2).

Considering the slow kinetics of oxime formation at neutral pH and the fact that catalysis-by-freezing cannot be applied for more sensitive proteins of interest, other means of catalysis were explored. A first option was to increase the salt concentration of the reaction mixture.^34^ It has been reported that an increased salt concentration can promote oxime formation, an effect ascribed to stabilization of the charged transition state in oxime formation which favors the rate limiting dehydration step.^34^ To evaluate the effects of an increased salt concentration, NaIO_4_-oxidized GBP-R86N was combined with a 50-fold molar excess of the biotin-PEG_3_-ONH_2_ label in a 0.1 M NaP_i_ buffer (pH 7) and increasing amounts of NaCl (0, 1, 2, or 3 M). Reactions were incubated at 20 °C and oxime formation was periodically monitored via LC-MS (Figure 4). While the control reaction in the absence of NaCl needed over 12h to reach completion, the addition of NaCl increased the reaction rate significantly. The reaction was completed within 8h in the samples containing 2 M and 3 M NaCl. This confirms that increasing the salt concentration can be an efficient way of catalyzing oxime ligation, at least for proteins that can remain in solution under high salt conditions.

**Figure 4.**
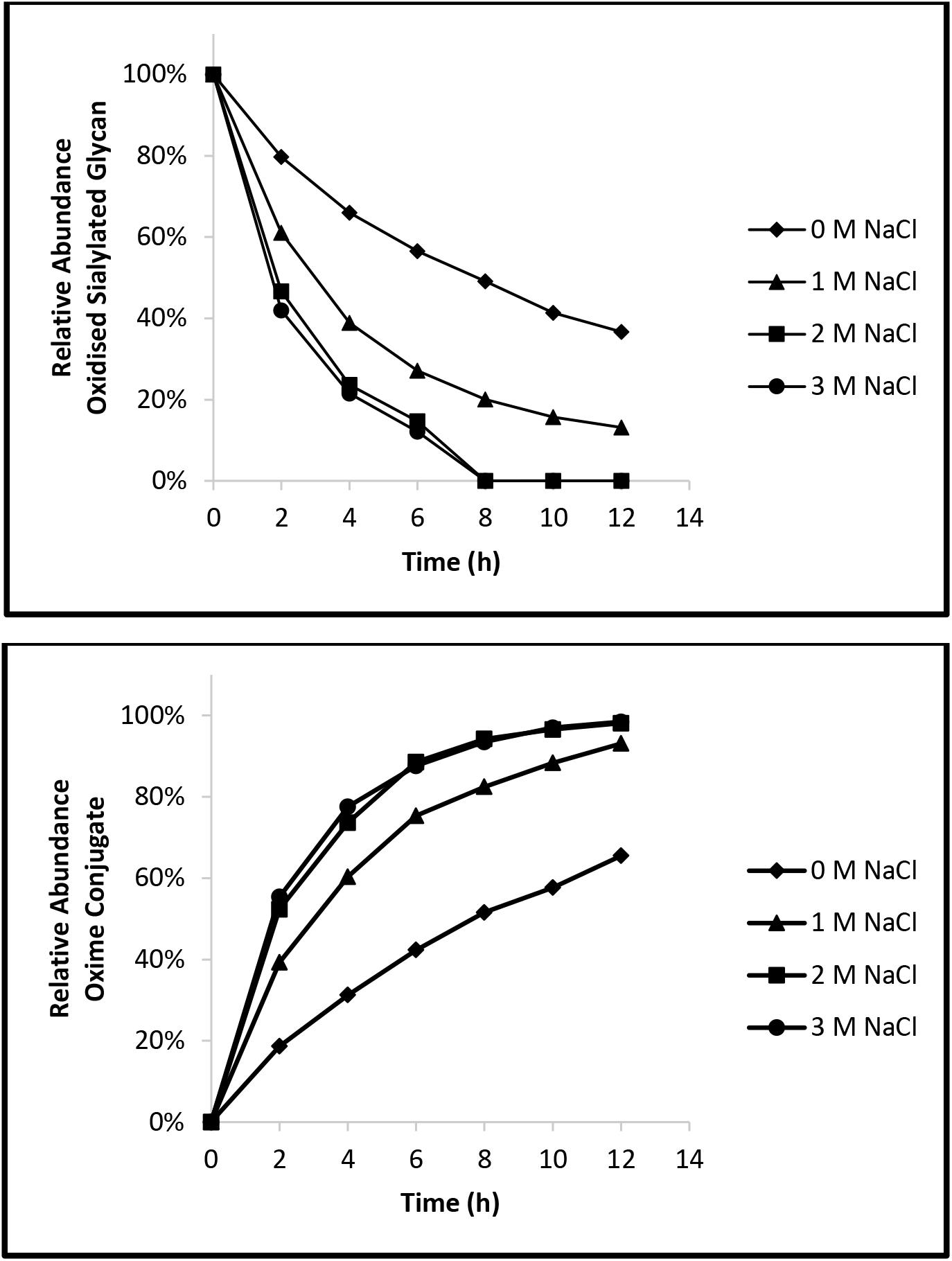
Effect of increased NaCl concentration on oxime ligation. NaIO_4_-oxidized GBP-R86N was conjugated with biotin-PEG3-ONH_2_ at increasing amounts of NaCl (0, 1, 2, or 3 M). Both the decrease of the oxidised LacNAc-Sia species (**top**) and the increase of the oxime conjugated product (**bottom**) were monitored over time via LC-MS. The relative abundance of the species of interest were plotted, using the abundance of the non-glycosylated GBP-R86N as an internal reference.

A second option for catalysis was the use of aromatic amines, previously described as efficient catalysts for oxime formation.^33,35–38^ NaIO_4_-oxidized GBP-R86N was mixed with a 50-fold molar excess of the biotin-PEG_3_-ONH_2_ label in a 0.1 M NaPi buffer (pH 7), either in the presence or absence of 50 mM aniline or PPD. Reaction mixtures were incubated at 20°C and analyzed via LC-MS analysis at regular time intervals to follow oxime formation (Supplementary Figure 4). As expected, uncatalyzed oxime formation progressed slowly and was still incomplete after 10h. Addition of either of the catalysts allowed complete and selective conjugation within 90 min. PPD-catalyzed conjugation was then further optimized, considering it was the most potent catalyst.^33,35^ Next, the effects of temperature on the PPD-catalyzed reaction was evaluated. Reaction mixtures containing NaIO_4_-oxidized GBP-R86N, 50 eq. of biotin-PEG_3_-ONH_2_ label and 50 mM PPD in a 0.1 M NaPi buffer (pH 7) were incubated at 4°C or 20°C and analyzed at regular time intervals via LC-MS (Supplementary Figure 5). At 20°C, PPD-catalyzed oxime formation was complete within 90 min. At 4°C, the reaction was considerably slower, yet still went to completion within 3.5h. Subsequent titration experiments showed that it is possible to reduce the PPD concentration to 1 mM and still obtain complete conjugation within 2h at 20 °C in the presence of 25 eq. of biotin-PEG_3_-ONH_2_.

As the oxime linkage is in principle a reversible bond that is subject to acid-catalysed dissolution, the stability of the oxime bond was assessed in the GlyConnect context. To this end, samples of biotin-PEG_3_-ONH_2_-conjugated GBP-R86N were exchanged to a pH 5 (0.1 M NaAc), pH 6 (0.1 M NaP_i_) or pH 7 (0.1 M NaP_i_) buffer. Vials were incubated at 37°C and sampled after 12h, 24h, 48h, 3 days and 3 weeks (Supplementary Figure 6). LC-MS analysis showed a steady decrease in the amounts of oxime conjugate at pH 5. In contrast, the analysis showed no or limited bond dissolution at pH 6 and pH 7, even after 3 weeks of incubation at 37°C, advocating that GlyConnect oximes are stable for longer time periods at near-neutral pH.

As the oxime ligation-based approach outlined above is dependent on the availability of an aminooxy-modified version of the label of interest, a limited commercial availability of these labels could hamper the applicability of the conjugation strategy. To expand the range of GlyConnect-compatible labels, we finally wanted to explore a reductive amination conjugation approach. In reductive amination, an aldehyde reacts with a primary amine-carrying label to form an imine, which is finally reduced to a secondary amine. Consequently, inclusion of reductive amination in the GlyConnect suite significantly expands the range of compatible and commercially available labels. To explore the reductive amination strategy, aniline was used as a label. As it was previously explored as a catalyst for oxime ligation, it was already established that aniline is reactive towards the aldehydes obtained through glycan oxidation and that the imine formed upon reaction with the aldehyde is unstable and short lived under the used aqueous conditions (addition of aniline was never observed in the oxime ligation experiments). Consequently, the presence of an aniline adduct provides adequate proof of successful reduction. As it is well established that a low pH promotes reductive amination,^39–42^ a range of different pH values were tested in the experimental setup. Upon NaIO_4_-oxidation of HEK293-GlycoDelete-produced GBP-R86N, using the standard conditions described above, protein samples were buffer exchanged to either a 0.1 M NaP_i_ buffer at pH 7 or pH 6, or to 0.1 M NaAc buffer at pH 5 or pH 4. Subsequently, a 50-fold molar excess of aniline was added to the samples, after which morpholine borane was added to a final concentration of 35 mM, and reaction mixtures were incubated for 4h at 20 °C. LC-MS analysis revealed a clear beneficial effect of low pH on reductive amination efficiency (Figure 5). At pH 5, the reductive amination had gone near to completion, whereas at pH 7 only a fraction of the available oxidized LacNAc-Sia had reacted. The data support the use of reductive amination as a valuable alternative to oxime ligation in the GlyConnect context.

**Figure 5.**
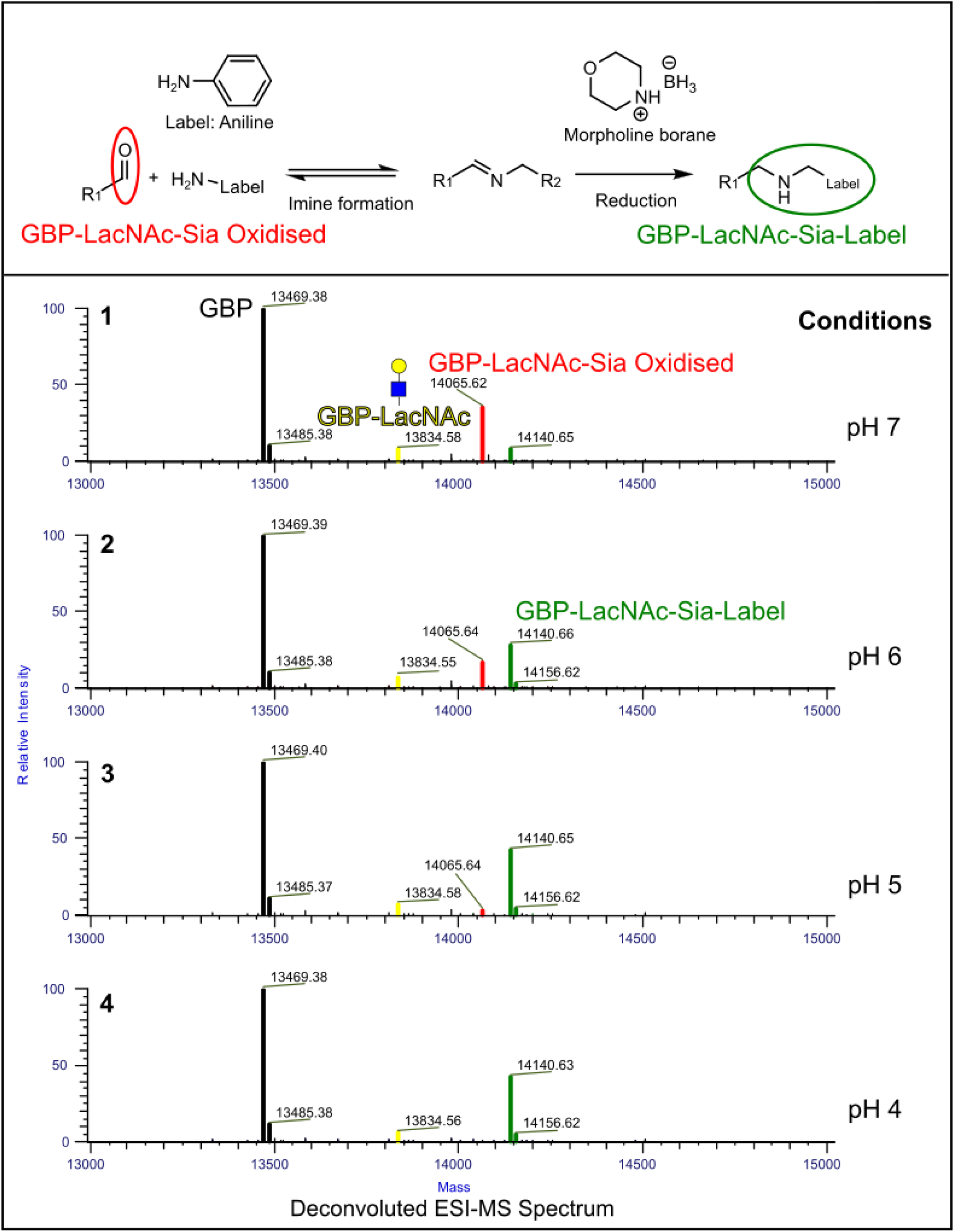
Conjugation of aniline on GlycoDelete-produced GBP-R86N through reductive amination. **Top.** Reaction scheme of the reductive amination with aniline. The aldehyde function (red) is obtained via NaIO_4_-based oxidation of the LacNAc-Sia glycan. First, the amine function of the aniline label reacts with the aldehyde to form an imine. Next, the imine double bound is reduced using morpholine borane. **Bottom.** The effect of pH on the morpholine borane based reductive amination conjugation. NaIO_4_ oxidized HEK293-GlycoDelete-produced GBP-R86N was used: a mixture of non-glycosylated VHH (black, MW = 13469 Da), LacNAc (yellow, MW = 13834 Da) and oxidized LacNAc-Sia species (red, MW = 14065 Da). The conjugated species signal appears at the expected mass (green, MW = 14140 Da).

### Engineering of the GlycoDelete strains for maximal GlyConnect compatibility

As shown in the previous section, efficient and selective conjugation can be reached using NaIO_4_ and GalOx based strategies. Importantly however, a matching glycosylation type is a prerequisite for efficient conjugation: the NaIO_4_-based method is dependent on the presence of the terminal Sia residue, while the GalOx based method requires a terminal Gal residue. To maximize GlycoDelete-GlyConnect compatibility, the mammalian and yeast GlycoDelete strains were further engineered to yield maximal amounts of the required glycans.

With the aim of obtaining increased amounts of sialylation, CRISPR/Cas9 was used with a tailored repair template to introduce the sialuria mutation in the UDP-N-acetylglucosamine-2-epimerase/N-acetylmannosamine kinase (GNE) gene of the HEK293-GlycoDelete strain. This mutation causes GNE to lose its sensitivity to feedback inhibition by CMP-sialic acid, and therefore leads to increased sialylation.^43,44^ Clonal selection and additional screening for high transfection efficiency and efficient growth yielded a performant HEK293-GlycoDelete-SiaHigh cell line. As expected, the increase in sialylation was found to be partially dependent on the target protein. GBP-R86N produced in this strain showed up to 30% increase in N-glycan sialylation compared to proteins produced in the parental HEK293-GlycoDelete strain (Figure 6).

**Figure 6.**
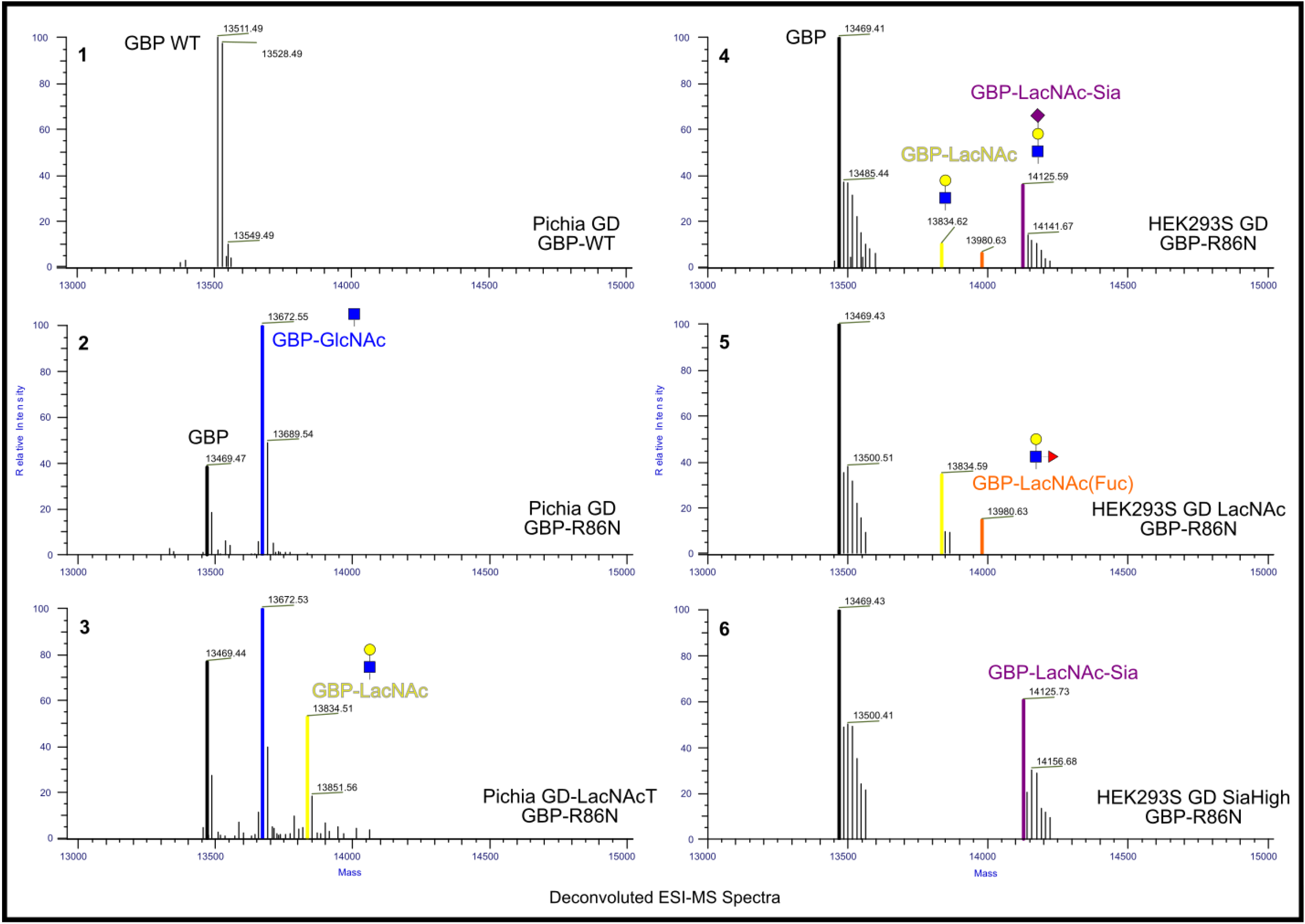
GlycoDelete strain optimization. GBP-(R86N) was produced in both Pichia-GlycoDelete strains (spectra 1-3) and in HEK293-GlycoDelete strains (spectra 4-6). In all spectra, both the pyroGlu and non-pyroGlu species are present. WT GBP (without glycosylation site; spectrum **1**) is not glycosylated, while GBP-R86N (with glycosylation site) is partially GlcNAc-modified (blue, MW = 13672 Da) (spectrum **2**). GBP-R86N produced in the Pichia-Glycodelete-LacNAc strain shows non-glycosylated (black, MW = 13469 Da), GlcNAc (blue, MW = 13672 Da) and LacNAc species (yellow, MW = 13834 Da) (spectrum **3**). GBP-R86N produced in the HEK293-GlycoDelete strain shows non-glycosylated (black, MW = 13469 Da), LacNAc (yellow, MW = 13834 Da), LacNAc(Fuc) (orange, MW = 13980 Da) and LacNAc-Sia species (purple, MW = 14125 Da) (spectrum **4**). Producing GBP-R86N in the HEK293-GlycoDelete-LAcNAc strain leads to the absence of the LacNAc-Sia species (spectrum **5**), while in the HEK293-GlycoDelete-SiaHigh strain the sialylation of the LacNAc glycan is complete (spectrum **6**).

To maximize LacNAc type glycosylation in the HEK293 background, a GNE gene knockout was generated in the HEK293-GlycoDelete line using CRISPR/Cas9.^43^ Subsequent screening and selection resulted in the HEK293-GlycoDelete-LacNAc cells. Several proteins were produced in this strain and all showed the presence of GlcNAc and LacNAc type GlycoDelete N-glycans, and the complete absence of sialylated N-glycans (LacNAc-Sia) (Figure 6).

In a parallel track, the *Pichia pastoris*-GlycoDelete strain was modified to allow in-cell synthesis of LacNAc N-glycans. This *Pichia Pastoris*-GlycoDelete-LacNAc strain was generated by introducing the human UDP-Glc-4-epimerase (GalE)^45^ gene and the human beta-(1,4) galactosyltransferase (GalT)^46^ gene in the *Pichia pastoris*-GlycoDelete strain. Up to 30% of GBP-R86N produced in this strain carried the LacNAc glycan (Figure 6).

### Expansion of the GlyConnect space

With a final set of proof-of-concept experiments, the technical feasibility of using the GlyConnect methodology for conjugation of other labels and proteins was tested.

A first experiment aimed at conjugating larger PEG polymers to GBP-R86N. Briefly, NaIO_4_-oxidized GBP-R86N in a 0.1 M NaPi buffer (pH 7) was mixed with varying amounts of aminooxy-functionalized PEG polymers of different chain lengths (5 kDa and 10 kDa; 25 or 10 eq.) and PPD (25, 5, or 1 mM). Reaction mixtures were incubated at 20 °C for 2h and subsequently analyzed via Western Blot (Figure 7) and LC-MS (Supplementary Figure 7). Analysis of the reaction mixtures showed successful glycan-specific conjugation of the longer PEG chains and pointed towards the combination of 1 mM PPD and 25 eq. of aminooxy-PEG polymer as the optimal condition within our test range. As these conditions are identical to the optimal conjugation conditions identified for the small biotin-PEG_3_-ONH_2_ label, it is fair to state that oxime reaction kinetics are not influenced by mere label size (at least within the scope of our experimental conditions). A follow-up experiment using the optimized conditions confirmed the feasibility of larger PEG chain conjugation, and even extended the size range to 20 kDa linear and Y-shaped PEG polymers (Supplementary Figure 8).

**Figure 7.**
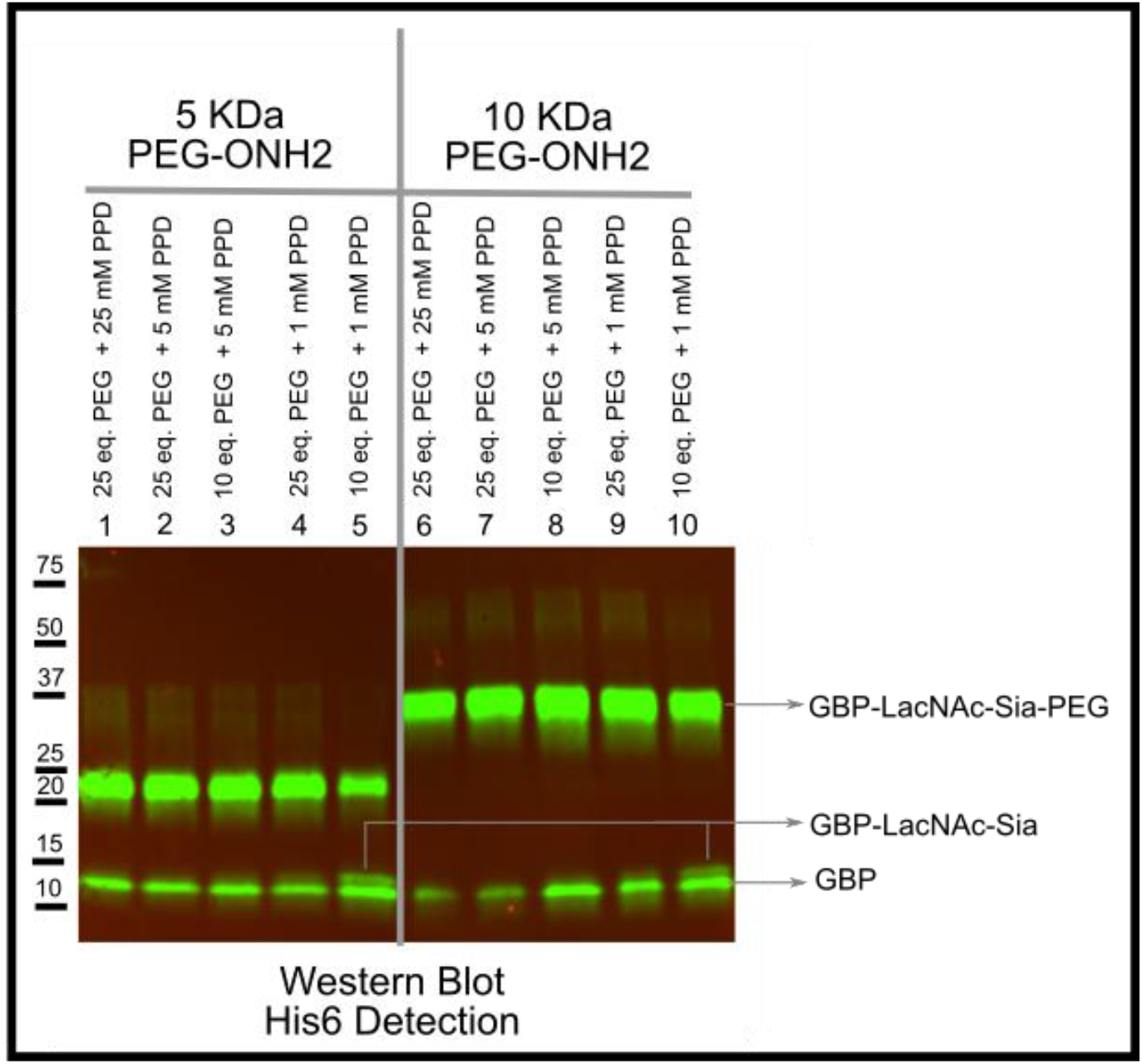
Western Blot analysis of oxime-based PEG polymer conjugation on GBP-R86N. NaIO4-oxidized HEK293-GlycoDelete-produced GBP-R86N, a mixture of non-glycosylated and LacNAc-Sia species, was conjugated with varying amounts of 5-kDa and 10-kDa aminooxy-PEG polymer, in the presence of varying concentrations of PPD.

In a second experiment, an amine-functionalized DTPA label was conjugated to GBP-R86N via reductive amination. Briefly, GBP-R86N was oxidized with 10 mM NaIO_4_ in NaP_i_ buffer (pH 7) as described before. After quenching the reaction, proteins were buffer exchanged to a 0.1 M NaAc buffer at pH 5, or the same buffer, containing 100 eq. of DTPA-NH_2_ and 35 mM of morpholine borane. Reaction mixtures were incubated overnight at 20 °C and subsequently analyzed via LC-MS. The results showed complete NaIO_4_ oxidation and almost complete DTPA conjugation of the sialylated species in the corresponding reaction mixtures, demonstrating that also this amine-functionalized chelator lends itself to efficient GlyConnect-based conjugation (Figure 8). The higher MW species detected at 53 Da above the expected DTPA-NH_2_ conjugate corresponds to the conjugated DTPA species chelated to a Ni(II)-ion (originating from the IMAC column during GBP-R86N purification).

**Figure 8.**
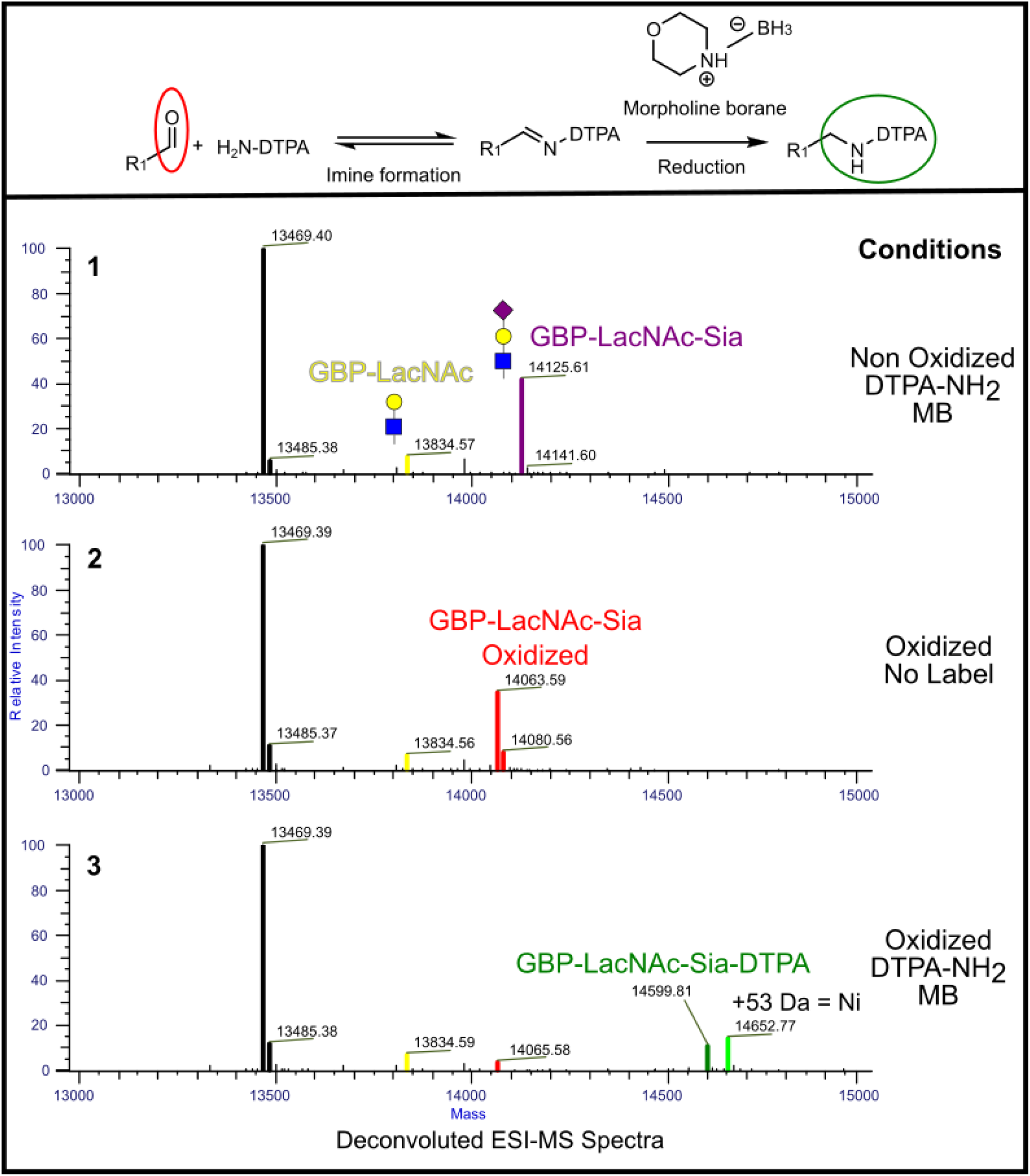
**Top.** Reaction scheme of the morpholine borane based reductive amination conjugation of amine functionalised DTPA label. The aldehyde function (red) is obtained via the selective oxidation of GlycoDelete glycans. **Bottom**. Deconvoluted MS spectra of the morpholine borane based reductive amination conjugation of amine functionalised DTPA on GBP-R86N. HEK293-GlycoDelete-produced GBP-R86N, a mixture of non-glycosylated (black, MW = 13469 Da), LacNAc (yellow, MW = 13834 Da) and LacNAc-Sia species (purple, MW = 14125 Da) (**1**), was oxidized (red, MW = 14063 Da) (**2**) after which DTPA-NH_2_ was conjugated through reductive amination (dark green, MW = 14599 Da) (**3**). Next to the DTPA-NH_2_ conjugated species, a +53 Da adduct appeared. This corresponds to the chelation of nickel to the DTPA conjugate (light green, MW = 14652 Da) (**3**).

In a third experiment, the GlyConnect methodology was tested on HEK293-GlycoDelete-produced hGM-CSF (with 6xHis tag). Protein samples were diluted in a 0.1 M NaP_i_ buffer (pH 7) and oxidized with 10 mM NaIO_4_. After buffer exchange to either a 0.1 M NaAc buffer (pH 5), a 0.1 M NaP_i_ buffer (pH 6), or a NaP_i_ buffer (pH 7) containing 1 mM PPD, we performed oxime ligation in the presence of 50 eq. of biotin-PEG_3_-ONH_2_ or 50 eq. of 10 kDa PEG-ONH_2_ polymer for 2h at 20 °C. Samples were analyzed via 6xHis- and biotin-specific Western Blotting (Figure 9). The oxidized protein samples showed successful conjugation, both with the small biotin-PEG_3_-ONH_2_ label (biotin detection) and the 10 kDa PEG-ONH_2_ polymer (6xHis tag detection). The observed heterogeneity of the conjugate signals can be explained by the fact that hGM-CSF has 2 N-linked glycosylation sites, in addition to several potential O-glycosylation sites. As both these glycan types can contribute to the degree of sialylation and as the nature of these glycans is inherently heterogeneous, this translates to a variable amount of conjugation sites per hGM-CSF molecule. This is reflected by an increased smearing of the signal for the small biotin-PEG_3_-ONH_2_ label and in a diffuse higher MW banded pattern for the larger 10 kDa PEG-ONH_2_ polymer. Non-oxidized samples showed only a band corresponding to unmodified hGM-CSF in the 6xHis-specific detection, evidencing conjugation specificity.

**Figure 9.**
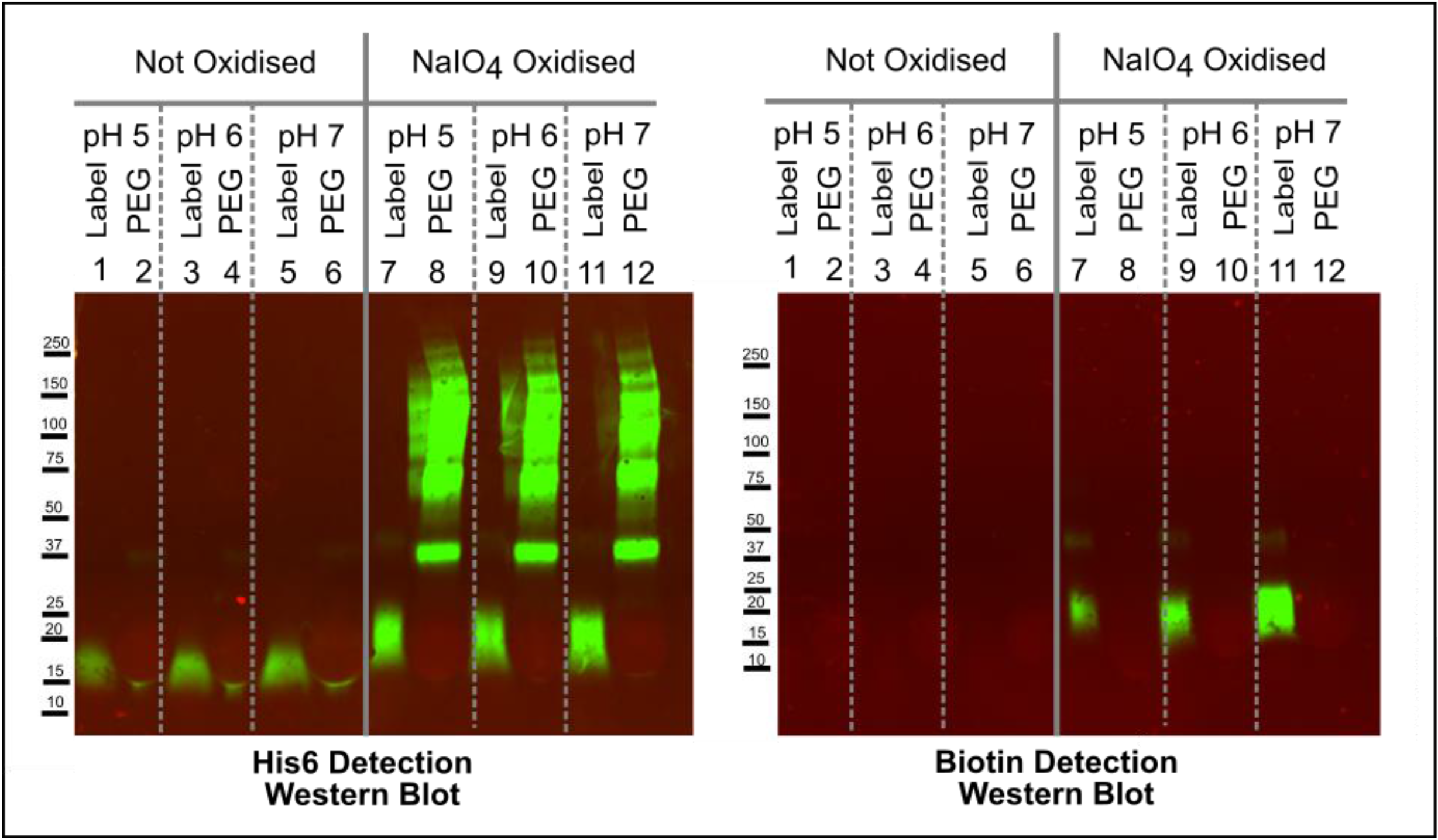
Western blot analysis of oxime-based PEG polymer conjugation on hGM-CSF. GlycoDelete-produced hGM-CSF was conjugated with either biotin-PEG_3_-ONH_2_ or 10 kDa PEG-ONH_2_ polymer at varying pH, with or without preceding oxidation. The left blot was visualized with a 6xHis tag specific fluorescent antibody, while the right blot was visualized with a biotin specific fluorescent antibody. hGM-CSF has multiple N-glycosylation sites (next to a number of O-glycosylation sites) leading to a higher degree of glycosylation and conjugation heterogeneity.

In a final experiment, the GlyConnect methodology was applied on a HEK293-GlycoDelete-produced obinutuzumab antibody. As preliminary tests had shown that NaIO_4_-based oxidation of the antibody glycan was ineffective at neutral pH, the pH of the 0.1 M NaP_i_ oxidation buffer was lowered to 6.5. Obinutuzumab samples were either left unoxidized, or were oxidized using 0.1 mM, 1 mM, 5 mM or 10 mM NaIO_4_. After buffer exchange to a 0.1 M NaP_i_ buffer (pH 7), oxime ligation was performed in the presence of 25 eq. of biotin-PEG_3_-ONH_2_ and 1 mM PPD for 2h at 20 °C. LC-MS analysis revealed that Sia oxidation was effective at 5 and 10 mM NaIO_4_, although some amino acid side oxidation was observed under these conditions (Figure 10). The subsequent conjugation reaction resulted in complete conjugation of the sialylated species.

**Figure 10.**
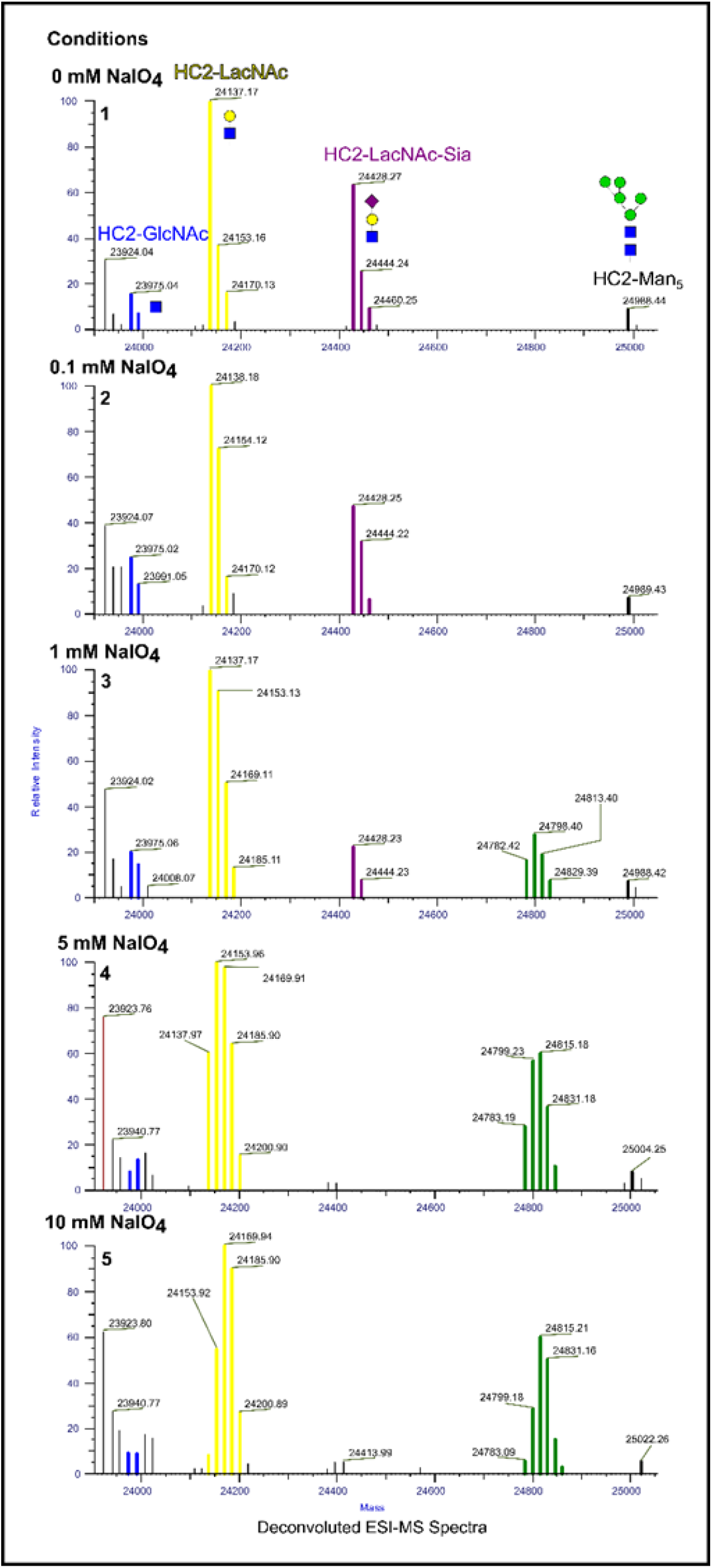
NaIO_4_-based oxidation and subsequent oxime conjugation of biotin-PEG3-ONH_2_ on obinutuzumab. HEK293-GlycoDelete-produced obinutuzumab, a mixture of GlcNAc (blue, MW = 23975 Da), LacNAc (yellow, MW = 24138 Da), LacNAc-Sia (purple, MW = 24430 Da) and GlcNAc_2_-Man_5_ species (black, MW = 24989 Da) (**1**), was oxidized at varying NaIO_4_ concentrations and subsequently conjugated with biotin-PEG3-ONH_2_ (green, MW = 24783 Da) (**2-5**).

## DISCUSSION

The aim of this study was to develop the GlyConnect platform as a reliable and broadly applicable conjugation extension of the previously developed GlycoDelete technology. GlycoDelete gives access to proteins with small and linear N-glycans that have either a terminal N-acetylglucosamine (GlcNAc), galactose (Gal) or sialic acid (Sia).^4^ The main pillar of GlyConnect is to use these unique glycans as handles for site-selective and homogeneous conjugation. In this way, the GlycoDelete/GlyConnect tandem opens up opportunities for applications in therapeutics, diagnostics, and the generation of research tools.

Five GlyConnect conjugation strategies were successfully explored on the glyco-engineered VHH GBP-R86N, produced in a GlycoDelete background. First, Sia-specific conjugation was performed using classical periodate oxidation in combination with oxime ligation. Second, Gal-specific conjugation was explored using GalOx-based oxidation in combination with oxime ligation. Third, another Gal-specific conjugation strategy was tested based on sialyltransferase-mediated derivatization of Gal with an azide-modified Sia, followed by strain-promoted alkyn-azide click chemistry. Fourth, GlcNAc-specific conjugation was tested through galactosyltransferase-mediated derivatization of GlcNAc with an azide-modified Gal (GalNAz), again followed by strain-promoted alkyn-azide click chemistry. Fifth, an alternative GlcNAc-specific conjugation was explored using a GlcNAc-reactive GalOx mutant (GalOx-F2) followed by oxime ligation. In these initial tests, the periodate-based strategy, the GalOx-based strategy and the GalNaz-click strategy went to completion, whereas the other strategies did not. GalOx- and GalOx-F2-based reactions showed evidence of non-specific oxidation. All other chemistries were proven to be highly specific. Only the periodate-based and GalOx-based methods were eventually selected for further optimization. Next to a high efficiency, these methods are established techniques in the glyco-engineering toolbox.^16,20,32,47^ Oxidation and conjugation reactions were optimized separately.

Periodate-based oxidation was optimized by varying the periodate concentration and pH of the oxidation buffer. The optimal condition was determined to be 10 mM NaIO_4_ in 0.1 M NaP_i_ (pH 7) (30 min at 4°C in the dark). Using these conditions, complete and Sia-specific oxidation of GBP-R86N was achieved. GalOx-based oxidation was optimized by varying temperature, reaction time, activity units, and the presence/absence of catalase and peroxidase. As GalOx reduces molecular oxygen to H_2_O_2_ during the oxidation of the primary alcohol on Gal, the H_2_O_2_ could in turn cause unwanted oxidation of the GalOx enzyme or the target protein. To avoid this, catalase can be added to the reaction mixture to convert the H_2_O_2_ into H_2_O and O_2_.^8,23,32,48^ Peroxidase can be added to the reaction mixture to re-activate the GalOx enzyme after oxidation of a substrate molecule.^48^ The optimization experiments showed that catalase and peroxidase were not required in the current context. Nevertheless, addition of these enzymes may be beneficial when working with other, more oxidation-sensitive target glycoproteins or in other experimental conditions. As there is a direct relationship between temperature and GalOx activity, but also an inverse relation between temperature and the solubility of O_2_ (a substrate of GalOx during the oxidation-reduction cycle) in water,^32,48^ it was imperative to scout for the optimal reaction temperature. Performing the reaction at 4°C allowed the complete oxidation of the target glycan, but was considerably slower than performed at 20°C. Titration experiments allowed determination of the optimal amount of GalOx units to be used for a fixed amount of target glycoprotein. The optimized conditions for LacNAc-specific GalOx-based glycoprotein oxidation were eventually determined to be 2U of GalOx for 1.6 nmol of protein (ca. 0.01 eq.) in a 0.1 M NaP_i_ (pH 7) (5h at 20°C).

As both the periodate- and the GalOx-based oxidation strategy yield reactive aldehydes, both strategies are compatible with the same conjugation chemistries. Oxime formation and reductive amination were our methods of choice. While oxime ligation results in an oxime bond that can be subject to acid-catalyzed hydrolysis, reductive amination yields a stable secondary amine bond which is much more stable.^22,42^ Different approaches to catalyze oxime formation were explored. Both catalysis-by-freezing at −20°C and performing the reaction in high salt promoted oxime formation, but obviously have a limited applicability for more challenging target proteins.^22,34^ As an alternative, aniline and p-phenylenediamine (PPD) were explored as organocatalysts for oxime formation, and PPD was identified as the most potent catalyst.^33,35^ The optimized condition was set at 1 mM PPD combined with 25 eq. of aminooxy label in a 0.1 M NaP_i_ (pH 7) (5h at 20°C). Of note is that the presence of PPD can inhibit GalOx activity, advocating a sequential oxidation and conjugation approach where PPD is only added to the reaction mixture after oxidation is complete.

Considering the high availability of amine-conjugates, reductive amination was evaluated as an alternative to oxime ligation. Reductive amination was optimized using aniline as a label and morpholine borane (MB) as a stable, water soluble reductive agent.^41,49,50^ A lower pH is known to be beneficial for reductive amination, as imine formation – the first step in the reaction mechanism – is acid-catalyzed.^39–41^ Therefore, the pH of the reductive amination reaction mixture was varied between pH 4 and 7. The optimized condition for reductive amination was 35 mM MB combined with 50 eq. of amine label in a 0.1 M NaAc buffer (pH 5) (4h at 20°C). Both the optimized oxime ligation and reductive amination strategies were found to give near-quantitative conjugation. The fact that oximes have been described as reversible bonds, the stability of the oxime linkage was evaluated at 37°C and at pH 5, pH 6 and pH 7. At pH 6 and 7, oxime hydrolysis was negligible after 72h. At pH 5, slow hydrolysis was observed. Given further optimization of the chemical environment of the reactive aminooxy group, oxime linkages could be exploited for pH-driven controlled release applications.^51,52^ Alternatively, if oxime hydrolysis is undesirable, a simple reduction of the oxime linkage can render it resistant to acid-catalyzed hydrolysis.^37,38,52,53^ Considering that the GlyConnect conjugation efficiency is intrinsically linked with the presence of a compatible glycan substrate, the GlycoDelete strains were further engineered to maximize GlyConnect compatibility. Because the periodate-based and the GalOx-based conjugation strategies were selected as the preferred conjugation methods, GlycoDelete strains were either engineered to increase the amounts of terminal sialylation (periodate-compatible) or terminal galactosylation (GalOx-compatible). For the mammalian GlycoDelete strains, engineering efforts centered around the UDP-N-acetylglucosamine-2-epimerase/N-acetylmannosamine kinase (GNE) enzyme, which is a key player in the synthesis of the CMP-sialic acid donor substrate for sialyltransferases.^43^ Introduction of the R263L sialuria mutation in the GNE gene of the HEK293-GlycoDelete strain yielded a HEK293-GlycoDelete-SiaHigh strain that exhibits increased N-glycan sialylation (i.e. higher LacNAc-Sia levels) as compared to the parental cells. The R263L mutation causes GNE to lose its sensitivity to feedback inhibition by CMP-sialic acid, resulting in increased CMP-sialic acid levels and increased sialylation in the cell.^43,44^ In contrast, a GNE knockout in the HEK293-GlycoDelete strain yielded a HEK293-GlycoDelete-LacNAc strain characterized by the absence of N-glycan sialylation and, consequently, increased amounts of terminal galactose (i.e. higher LacNAc levels). GNE knockout impairs sialylation by precluding synthesis of the CMP-sialic acid donor substrate. In a parallel track, the *Pichia pastoris*-GlycoDelete strain was modified to allow in-cell synthesis of LacNAc N-glycans. Stable expression of human UDP-Glc-4-epimerase (GalE; converts UDP-Glc to UDP-Gal) and human beta-(1,4) galactosyltransferase (GalT; uses UDP-Gal as a substrate and transfers Gal to GlcNAc, yielding LacNAc) in the *Pichia pastoris*-GlycoDelete background yielded a *Pichia Pastoris*-GlycoDelete-LacNAc strain. Initial transformations of this strain with GBP-R86N yielded protein with up to 30% LacNAc type glycosylation. Undoubtedly, further engineering, clone selection and optimization of culture conditions will lead to a further increase in protein LacNAc glycosylation.

Finally, to estimate the value and technical feasibility of the GlyConnect strategy, the optimized chemistries were now tested with different labels and target proteins. Successful conjugation of larger PEG polymers (ranging from 5 kDa to 20 kDa) on GBP-R86N showed that the optimized conjugation conditions are likely independent of label size. Also, reductive amination-based conjugation of an amine-functionalised DTPA (a chelator used in medical radio-imaging)^54–56^ on GBP-R86N proved to be successful. In another example, successful oxime-based conjugation of both short and longer PEG polymers to GlycoDelete-produced human granulocyte-macrophage colony-stimulating factor (hGM-CSF) was achieved. GlyConnect-based conjugation of PEG or other half-life extending moieties to this molecule could be useful in a next-generation therapeutic hGM-CSF.^57,58^ A final example involved the oxime-based conjugation of a small aminooxy label to a GlycoDelete-produced, full-length human IgG1 antibody (obinutuzumab). While our standard conditions for sialic acid-specific oxidation were proven to be ineffective on the antibody, a slight shift to a more acidic pH (6.5) was enough to achieve complete periodate-based oxidation of the sialylated species.^17^ However, under these conditions, some off-target amino acid oxidation was observed (most likely methionine). Attempts to modify the antibody via a GalOx-based methodology failed, possibly due to steric hindrance (data not shown).

Countless applications in life sciences depend on the availability of complex natural and recombinant (glyco)proteins.^59,60^ Conjugation of these proteins to other molecules allows for an additional layer of complexity and control, and often greatly expands their applicability.^61–64^ It is therefore not surprising that the past decades have seen the advent of several (semi-) site-specific protein conjugation strategies, which can broadly be divided into natural amino acid-directed, unnatural amino acid-directed, and glycan-directed chemistries.^16,63,65–69^

The most commonly used protein conjugation strategies are, arguably, the classical chemistries directed against natural amino acids. Typical examples include Lys- and Cys-specific chemistries, and chemistries that target the N- or C-terminus of the protein of interest.^65,69,70^ Obvious advantages of these strategies include that they can generally be performed on native, non-engineered proteins, and that an ample range of labels/cargo molecules with suitable functional groups is readily available. A notable drawback these strategies have in common is the limited control over the label:protein ratio, as the frequency of occurrence of specific amino acid residues is hard-coded into the protein’s primary amino acid sequence (e.g. more than 20 surface-exposed Lys and 8 hinge-site Cys residues present in an average human IgG1 antibody)^16^ and accessibility of a residue for conjugation is governed by the protein’s secondary and tertiary structure.^71^ Other difficulties relate to specificity (e.g. Lysine vs N-terminal amino acid), negative effects on protein structure (e.g. need for reduction of disulfide bridges to allow thiol coupling and potential disulfide scrambling), and the frequent loss in conjugation efficacy when trying to mitigate aforementioned hurdles.^64,72^ More recently, methodologies harnessing the power of specific enzymes like Sortase, Transglutaminase or Formylglycine-Generating enzyme have expanded the toolbox for natural amino acid-directed conjugation.^29,70,73–77^ These chemo-enzymatic strategies allow more control over label:protein ratio and site-specificity as they target specific AA stretches or so-called ‘tags’. On the other hand, their very dependency on tags also confines their use, as the protein of interest must tolerate the introduction of a specific consensus sequence without repercussions on protein structure and activity.

The most recent development in the protein conjugation field is the use of unnatural amino acids (UAAs).^67,68^ Conjugation technology based on UAAs overcomes the major hurdles encountered with natural amino acid-directed conjugation. The UAA can in fact be seen as a one-amino-acid tag that introduces a unique bio-orthogonal chemical handle on a well-defined site in the protein, allowing for high-yielding and site-specific conjugation. The flip side of the coin is however that the use of UAAs requires extensive expression platform engineering, protein engineering and expensive tailored growth media, arguably making the process more cumbersome and less scalable. ^8,67,71,78,79^

Glycan-based conjugation sets itself apart from the natural amino acid- and UAA-based conjugation approaches by the fact that a cargo gets linked to the protein through a glycan, which acts as a flexible linker.^46,61^ It lends itself most obviously for conjugation of proteins that are glycosylated in their native form, although proteins that are not natively glycosylated can be engineered to contain an N-x-T/S N-glycosylation signature. Glycan-based coupling can be a powerful strategy for glycoprotein conjugation, as the glycans on a glycoprotein are generally quite accessible for conjugation, and aid in pinpointing the sites where bulky groups can be tolerated. Importantly, glycan-based coupling allows chemical or chemo-enzymatic glycoprotein labelling without a direct modification on the protein itself, thus lowering the risk of disrupting the protein structure and function.^8,14,80^ Of note is that not only N-glycans, but also O-glycans can be derivatized via glycan-targeted conjugation, provided they present suitable residues at their outer termini. The classical glycan-directed conjugation approaches selectively oxidize the glycans to generate reactive aldehydes, which are then reacted with compatible functional groups. A notorious difficulty with these conjugations lies in the heterogeneous nature of glycans,^2,6^ which can translate to variability in conjugate structure and conjugation numbers.^80,81^ An alternative and more recent methodology for glycan-directed conjugation consists of a multistep approach, wherein the native N-glycans are partially trimmed and subsequently derivatized with a bio-orthogonal function.^8,30^ Finally, the bio-orthogonal function can be used to couple a molecule of interest. A popular format is the grafting of an azide-functionalized monosaccharide onto a trimmed glycan. Commercial kits are available for this, of which the GlyClick Antibody Labelling kit used in this study is an example. Although these strategies allow site-selective and homogeneous conjugation, processing is quite cumbersome and scalability may be an issue. The GlycoDelete/GlyConnect tandem described in this study expands the glycan-directed conjugation toolbox and reduces some of the biggest hurdles of glycan-based conjugation. The GlycoDelete technology allows a standard protein production workflow in standard growth media and provides immediate access to short and linear glycans that act as linkers and anchor points for protein coupling. Though GlycoDelete-type glycosylation may also be obtained through enzymatic de- and re-glycosylation, the *in cellulo* approach that forms the basis of the GlycoDelete technology obviates the need for cumbersome *in vitro* pre-processing steps. The GlyConnect arm allows efficient conjugation to the glycan linker, either through classical oxidation-based approaches or through the azide-alkyne click methodology described above. In sum, the GlycoDelete/GlyConnect methodology presented here boasts a unique set of features that can make it the method of choice for a broad range of protein conjugation needs.

## AUTHOR CONTRIBUTIONS

WVB and KT designed and performed all conjugation experiments. WVB, FS, SV, EW generated the HEK293-GlycoDelete-SiaHigh and HEK293-GlycoDelete-LacNAc strains, and BL, BVM, SV generated the Pichia-GlycoDelete-LacNAc strain. GalOx was produced and purified by WVB, BL and SV. GlycoDelete-proteins were produced and purified by SV and EW. SD and FS developed the LC-MS methods. WN provided advice on conjugation chemistry. WVB, KT and SD wrote the manuscript. AM and NC supervised the work.

