## Supplementary Information for "GlyConnect: a glycan-based conjugation extension of the GlycoDelete technology" (from another version of this preprint: version 1, 2021-06-02)

#### Production of galactose oxidase

An expression vector was made wherein the coding sequence of the GalOx enzyme of the fungus *Fusarium graminearum* is operably linked to the methanol-inducible AOX1 promoter of *Pichia pastoris*. The CDS encodes (from N- to C-terminus) the *S. cerevisiae* Mating Factor pre-propeptide, a GS linker (GS), amino acid 42 to 680 of the *F. graminareum* GalOx, a GS linker (GS), and an 8xHis tag. The resulting expression vector was introduced into the Pichia pastoris-GlycoDelete strain, and transformants were plated on selective medium for clone selection. For enzyme production, a selected transformed *Pichia pastoris* clone was first grown for 48h at 28°C in BMGY medium (100 mM potassium phosphate pH 6, 2%$_{w/v}$ peptone, 1%$_{w/v}$ yeast extract, 1%$_{w/v}$ yeast nitrogen base without amino acids and 1% glycerol) containing glycerol as the sole carbon source; this starter culture was then used to inoculate 1L of BMMY (100 mM potassium phosphate pH 6, 2%$_{w/v}$ peptone, 1%$_{w/v}$ yeast extract, 1%$_{w/v}$ yeast nitrogen base without amino acids and 1% methanol; induction of recombinant protein expression by substitution of glycerol for methanol) containing 0.5mM $CuSO_4$ at an OD600 of 1. The culture was maintained for 96h at 28°C, with the addition of 1% methanol every 24h, upon which the growth medium (supernatant) of the culture was collected via centrifugation (1500xg, 4°C, 10 minutes). Supernatant was filtered using a 0.22µm filter, upon which 100 mg/L reduced glutathione and 2 mM $MgSO_4$ were added, the pH was adjusted to pH 7.5, and precipitates were cleared using a 0.22µm filter. Finally, enzyme was purified from the cleared supernatants via consecutive IMAC (HiTrap IMAC HP 5 ml, GE Healthcare) and SEC (HiLoad Superdex 200 pg, GE Healthcare; PBS as running buffer) chromatography using standard protocols. Purified enzyme was aliquoted, snap-frozen in liquid nitrogen, and stored at -70°C until use. The specific activity of GalOx stocks was assayed with the Amplex® Red Galactose/Galactose Oxidase Kit (Thermo Fisher Scientific™) using Gal as a substrate, following the manufacturer's instructions.

#### Production of benchmark GlycoDelete proteins

To generate HEK293-GlycoDelete-produced, HEK293-GlycoDelete-LacNAc-produced and HEK293-GlycoDelete-HighSia-produced GBP-R86N, the respective cell lines were transfected with a pcDNA3.3TOPO expression vector encoding the glyco-engineered VHH GBP-R86N-His6

(*MGWSCIIFFLVATATGVHS*
QVQLVESGGALVQPGGSLRLSCAASGFPVNRYSMRWYRQAPGKEREWVAGMSSAGDRSSYEDSVKGRFTISRDDANNTVYLQMNSLKPEDTAVYYCNVNVGFEYWGQGTQVTVSSHHHHHH; Mouse Ig Heavy Chain signal sequence in italics) using PEI as described before. Transfected cells were kept in a 1:1 mixture of Freestyle293 medium and Ex-Cell293 medium (FREX293 medium) for 72h at 37°C, after which the culture medium was collected. Recombinant protein was purified from the culture medium via consecutive immobilized metal affinity chromatography (IMAC; $Ni^{2+}$-loaded HisTrap) and size exclusion chromatography (SEC; Superdex 75; PBS as running buffer) purification. The purity and composition of each protein batch was confirmed using Coomassie Blue analysis, 6xHis-tag-specific Western Blot analysis and LC-MS. Proteins were snap frozen and kept at -80°C until use.

To generate HEK293-GlycoDelete-produced hGM-CSF, HEK293-GlycoDelete cells were transfected with the pORF-hGM-CSF-6xHis vector, encoding hGM-CSF-6xHis (*MWLQSLLLLGTVACSIS*
APARSPSPSTQPWEHVNAIQEARRLLNLSRDTAAEMNETVEVISEMFDLQEPTCLQTRLELYKQGLRGSLTKLKGPLTMMASHYKQHCPPTPETSCATQTITFESFKENLKDFLLVIPFDCWEPVQEHHHHHH; native signal sequence in italics) under control of a hEF1a/HTLV1 promoter. Protein production, purification and quality control were performed as described for HEK293-GlycoDelete-produced GBP-R86N-6xHis.

To generate HEK293GlycoDelete-derived obinutuzumab antibody, HEK293-GlycoDelete cells were transfected with a dual expression vector, encoding the obinutuzumab heavy chain in one CMV-controlled expression cassette and the obinutuzumab light chain in a second, separate CMV-controlled expression cassette *(expression vector obtained from NovImmune)*. Protein production was performed as described for HEK293GlycoDelete-derived GBP-R86N-His6 protein. The recombinant obinutuzumab protein was purified from the culture medium through consecutive IMAC ($Ni^{2+}$-loaded HisTrap) and SEC (Superdex 200; PBS as running buffer) purification. The purity and composition of the protein was confirmed using Coomassie Blue analysis, human IgG-specific Western Blot analysis and LC-MS. Proteins were snap frozen and kept at -80°C until use.

To generate Pichia-GlycoDelete-produced GBP-R86N and Pichia-GlycoDelete-LacNAc-produced GBP-R86N, the respective Pichia strains were transformed with a plasmid encoding the glyco-engineered GBP-R86N-8xHis benchmark protein

(*MRFPSIFTAVLFAASSALAAPVNTTTEDETAQIPAEAVIGYSDLEGDFDVAVLPFSNSTNNGLLFINTTIASI AAKEEGVSLEKREAEA* GSQVQLVESGGALVQPGGSLRLSCAASGFPVNRYSMRWYRQAPGKEREWVAGMSSAGDRSSYEDSVK GRFTISRDDANNTVYLQMNSLKPEDTAVYYCNVNVGFEYWGQGTQVTVSSGSHHHHHHHH; S. cerevisiae Mating Factor pre-propeptide in italics) under control of the methanol inducible AOX1 promoter. Transformed P. pastoris cultures were grown for 48h at 28°C in BMGY medium (100 mM potassium phosphate pH 6, $2\%_{w/v}$ peptone, $1\%_{w/v}$ yeast extract, $1\%_{w/v}$ yeast nitrogen base without amino acids and 1% glycerol) containing glycerol as the sole carbon source, and subsequently transferred to BMMY medium (100 mM potassium phosphate pH 6, $2\%_{w/v}$ peptone, $1\%_{w/v}$ yeast extract, $1\%_{w/v}$ yeast nitrogen base without amino acids and 1% methanol) which induced recombinant protein expression by substitution of glycerol for methanol. After 48 hours of incubation at 28°C during which 1% methanol was spiked three times, growth medium (supernatant) of each recombinant culture was collected. The recombinant GBP-R86N-8xHis proteins were purified via IMAC ($Ni^{2+}$-loaded HisTrap) and size exclusion chromatography (Superdex 75; PBS as running buffer). The purity and composition of each protein batch was confirmed using Coomassie Blue analysis, 6xHis-tag-specific Western Blot analysis and LC-MS. Proteins were snap frozen and kept at -80°C until use.

**GlycoDelete platform engineering**

To generate the HEK293-GlycoDelete-SiaHigh strain, the HEK293-GlycoDelete cell line was modified using GNE-specific CRISPR-Cas9 editing in combination with a repair template. The guide and the repair template were designed using Benchling (https://www.benchling.com). A GNE_guideE5fw (5'-cacc**G**AAGAGATGGTTCGAGTGATG-3'; overhang in lowercase; **G** added for U6-promoter compatibility) and a GNE_guideE5rev (5'-aaacCATCACTCGAACCATCTCTT**C**-3'; overhang in lowercase; **C** added as complement for the **G**) DNA oligo were heteroduplexed and cloned into a BbsI-digested pSpCas9(BB)-2A-GFP vector (Addgene) as described by Ran et al., 2013[24]. The repair template (5'-TTTACGTGGGCTATACTTGCCAATGAAAATGGTTTCTTTATCTCACAAACCATTTACCAATTGGTTCTT TTCATTTTCATAGGGAGCAAAGAGATGGTT**cta**GTGATG*aga*AAGAAGGGCATTGAGCATCATCCCAA CTTTCGTGCAGTTAAACACGTCCCATTTGACCAGTTTATACAGTTGGTTGCCCATGCTGGCTG-3') was ordered as a gBlock (IDT) and contained the R263L sialuria mutation (CGA→CTA; lowercase bold) and a CGG→AGA mutation (lowercase italics) to remove the PAM sequence. The vector co-expressing the guide RNA, Cas9 and GFP was co-transfected with the repair template into HEK293GlycoDelete cells using FuGENE® 6 Transfection Reagent (Promega). At 48h post transfection, cells were FACS-sorted and pooled based on GFP expression, and the GFP expressing pool was kept in culture for 48h. Finally, single clones were obtained using FACS sorting and clonal expansion was performed over a period of 3 weeks. Clones were then genotyped via PCR, restriction digest (the edited sequence is sensitive to BfaI digest, the parental sequence is not; the parental sequence is sensitive to TaqαI digest, the edited sequence is not) and Sanger sequencing, and homozygous edited clones were selected. Upon further selection for growth and transgene expression rates, a final clone was selected as the HEK293-GlycoDelete-HighSia line and cell-banked. To confirm the phenotype, cells were transfected with a pcDNA3.3TOPO expression vector (Thermo Fisher Scientific™) encoding the glyco-engineered benchmark VHH GBP-R86N-6xHis (preceded by an in-frame Mouse Ig Heavy Chain signal sequence), and the produced VHH was analyzed via LC-MS.

To generate the HEK293-GlycoDelete-LacNAc strain, the HEK293-GlycoDelete strain was modified using GNE-specific CRISPR-Cas9 editing. Upon design of the guide with Benchling, a GNE_E2_G3_FWD (5'-cacc**G**TTTAATGCCAAACATGATCG-3'; overhang in lowercase; **G** added for U6-promoter compatibility) and a GNE_E2_G3_REV (5'-aaacCGATCATGTTTGGCATTAAA**C**-3'; overhang in lowercase; **C** added as complement for the **G**) DNA oligo were heteroduplexed and cloned into a BbsI-digested pSpCas9(BB)-2A-GFP vector as described before. The vector co-expressing guide RNA, Cas9 and GFP was transfected into HEK293-GlycoDelete cells using FuGENE®. At 48h post transfection, cells were FACS-sorted and pooled based on GFP expression, and the GFP expressing pool was kept in culture for 48h. Finally, single clones were obtained using FACS sorting and clonal expansion was performed over a period of 3 weeks. Clones were then genotyped via PCR and Sanger sequencing, and knockout clones were selected. Upon further selection for growth and transgene expression rates, a final clone was selected as the HEK293-GlycoDelete-LacNAc strain and cell-banked. To confirm the phenotype, cells were transfected with a pcDNA3.3TOPO expression vector encoding GBP-R86N-6xHis (preceded by an in-frame Mouse Ig Heavy Chain signal sequence), and the produced VHH was analyzed via LC-MS.

To generate the Pichia-GlycoDelete-LacNAc strain, the Pichia-GlycoDelete strain (unpublished results) was transformed with a linearized pGAPNORMnn2SpGal10hGalT-I GlycoSwitch vector.[25] This vector encodes a fusion protein consisting of the N-terminal Golgi-localization domain of the S. cerevisiae Alpha-1,2-mannosyltransferase (MNN2), in frame with the entire *Schizosaccharomyces pombe* UDP-Gal 4-epimerase and the C-terminal catalytic domain of human β-1,4-galactosyltransferase 1 (B4GALT1). Transformants were grown on selective medium, and several clones were picked. Clones were phenotyped by subsequently transforming them with a vector encoding a glyco-engineered benchmark VHH GBP-R86N-6xHis under control of the AOX1 promoter. The produced VHH was analyzed with LC-MS. Based on the VHH glycoprofile and recombinant protein yields, one clone was selected as the Pichia-GlycoDelete-LacNAc strain and cell-banked.

#### Supplementary Figures

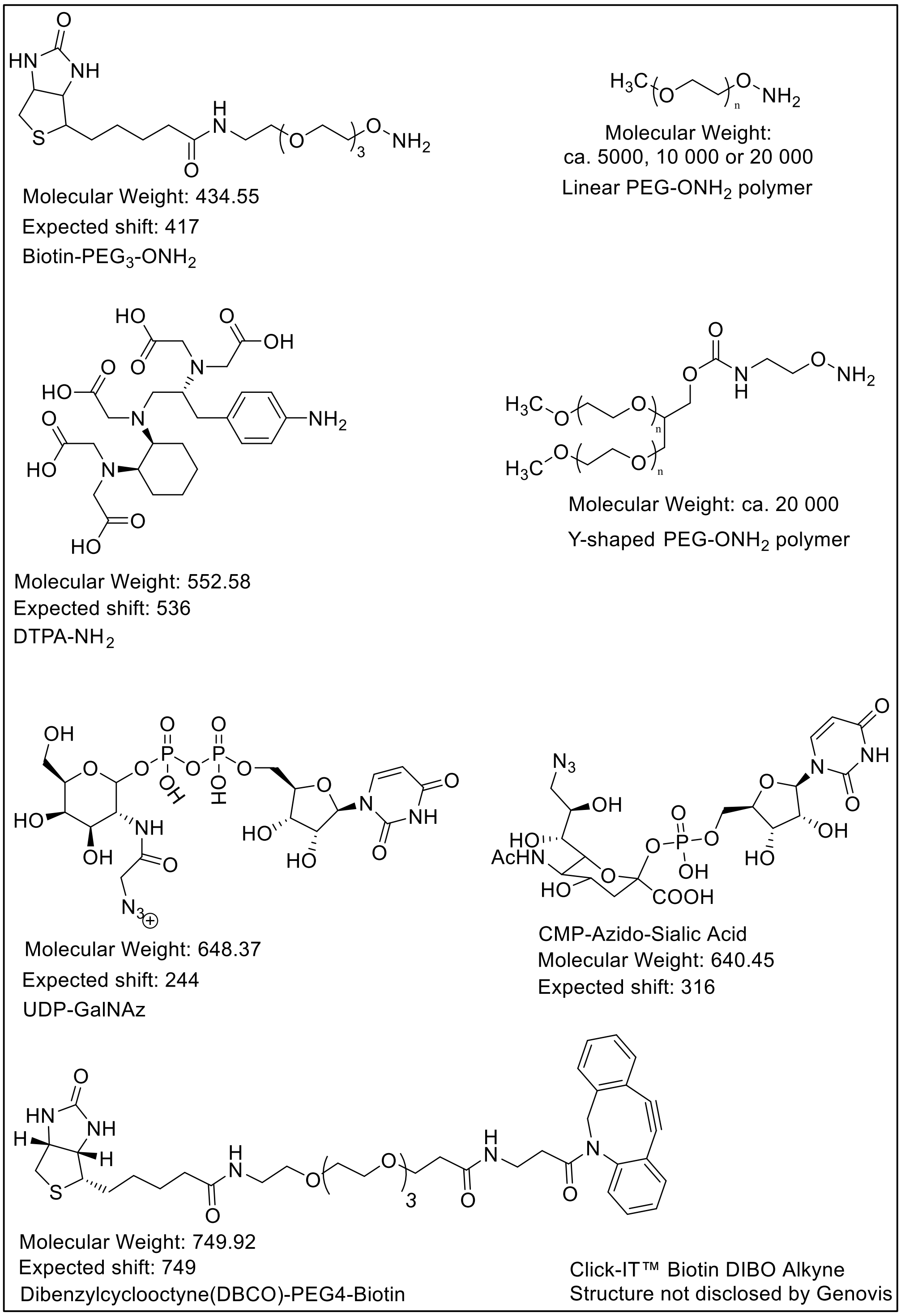


**Supplementary Figure 1.** Label structures.

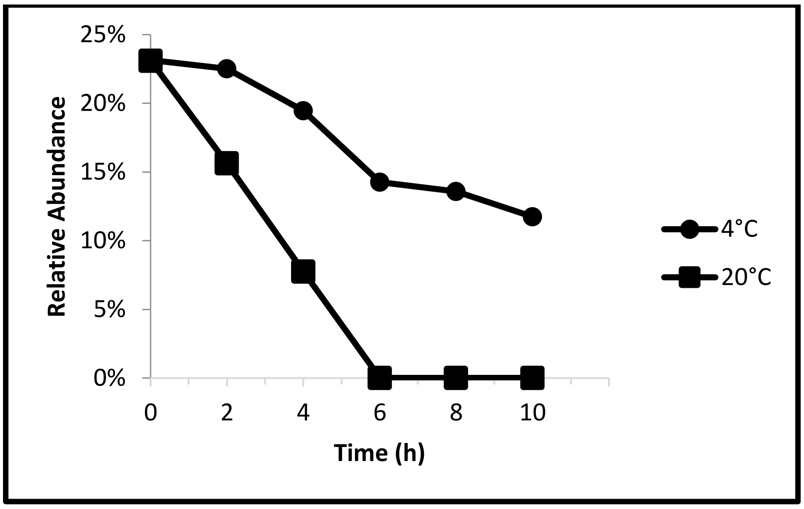


**Supplementary Figure 2.** Decrease in relative abundance of the non-oxidized LacNAc fraction of the GalOx based oxidation in function of temperature, as monitored with LC-MS. The relative abundance of the species of interest were plotted, using the abundance of the non-glycosylated GBP, which does not participate in the reactions, as internal reference.

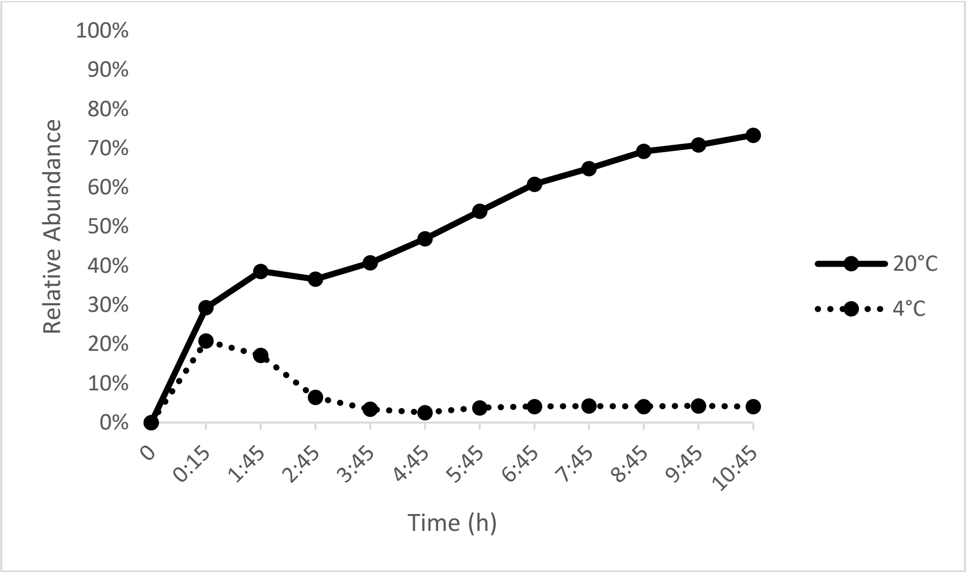


**Supplementary Figure 3.** Effect of temperature on the oxime bond formation towards $NaIO_4$ oxidized HEK293-GlycoDelete-produced GBP-R86N. Oxime conjugations were performed at 4°C and 20°C with biotin-$PEG_3$-$ONH_2$ label, and the increase in oxime conjugation was followed in function of time, via LC-MS. The relative abundances in the deconvoluted MS spectra, of the species of interest, were plotted using the abundance of the non-glycosylated GBP, which does not participate in the reactions, as internal reference.

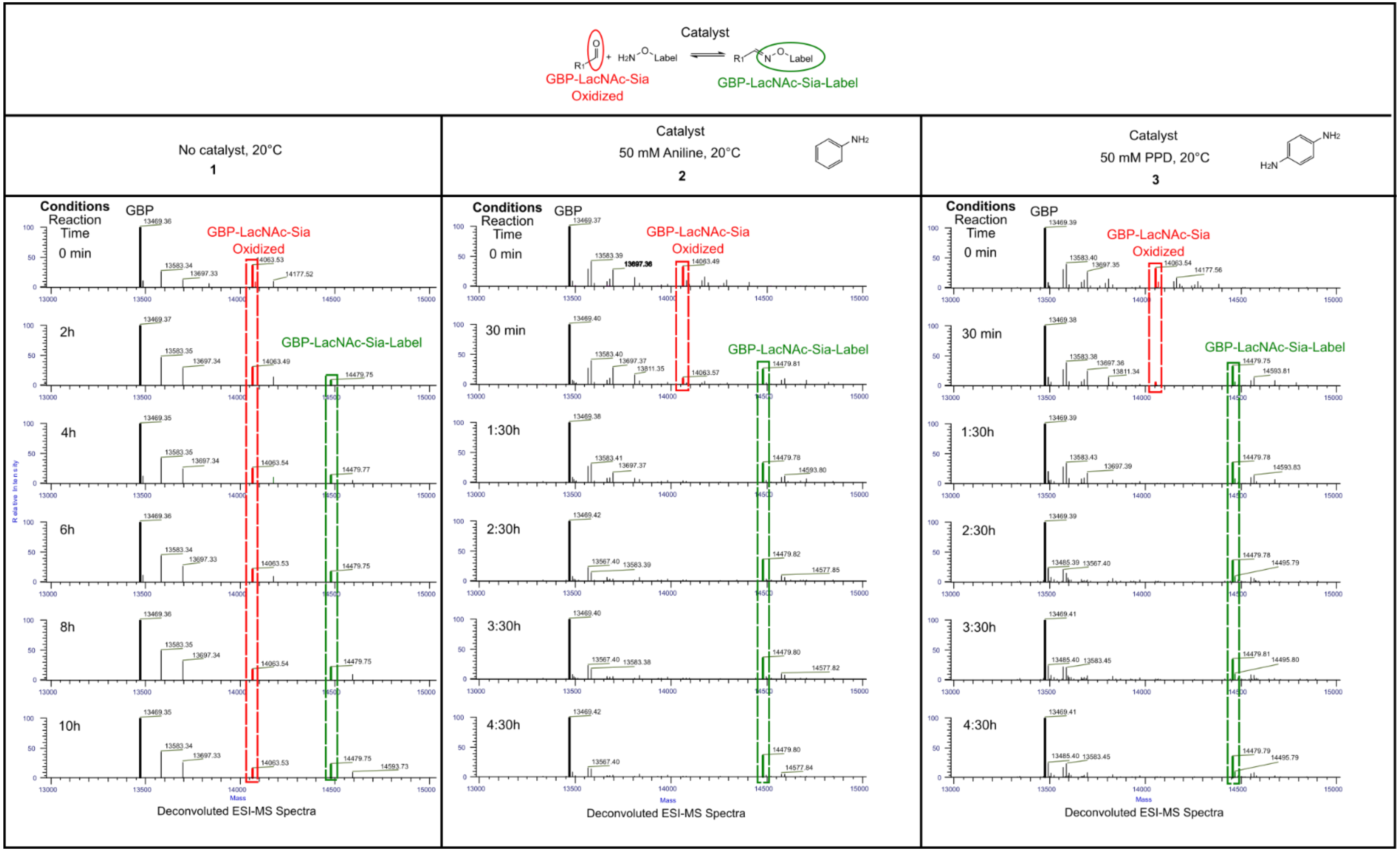
Catalyst
GBP-LacNAc-Sia Oxidized
GBP-LacNAc-Sia-Label
No catalyst, 20°C
1
Catalyst
50 mM Aniline, 20°C
2
Catalyst
50 mM PPD, 20°C
3
Conditions Reaction Time
GBP
Deconvoluted ESI-MS Spectra

**Supplementary Figure 4. Top**. Reaction scheme of the amine catalysed oxime formation. The aldehyde (red) originates from an oxidised LacNAc-Sia glycan ($R_1$) and is reacted with an aminooxy functionalised label in the presence of a catalyst, to form the oxime conjugate (green). **Bottom.** The effect of different amine catalysts on oxime conjugation. Oxidized HEK293-GlycoDelete-produced GBP-R86N, a mixture of non-glycosylated (black, MW = 13469 Da) and LacNAc-Sia species (red, MW = 14063 Da), was ligated with biotin-$PEG_3$-$ONH_2$ in the presence of different catalysts. The non-glycosylated species serves as internal control. The conjugated species signal appears at the expected mass (green, MW = 14479 Da).

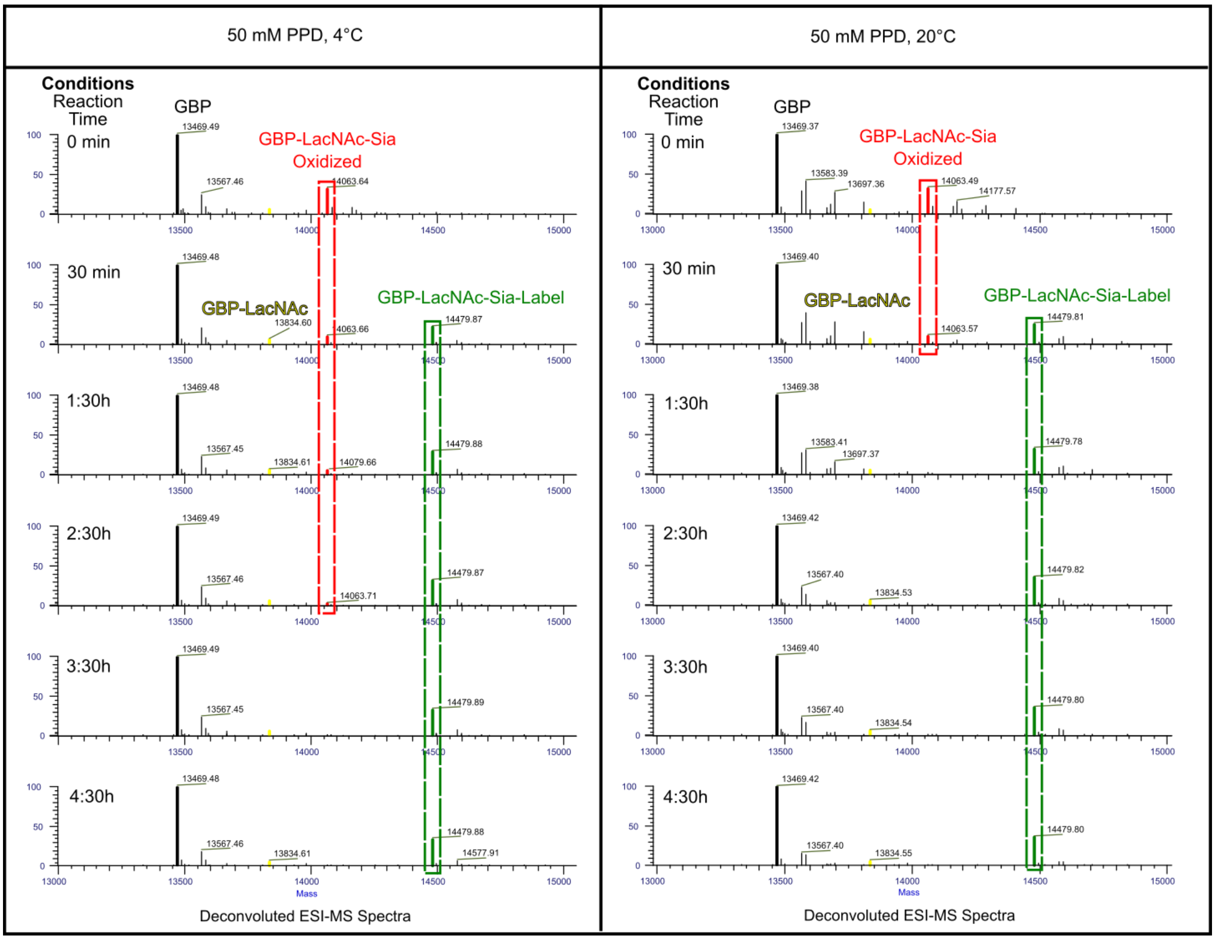


**Supplementary Figure 5**. The effect of temperature on PPD-catalysed oxime conjugation of biotin-PEG$_3$-ONH$_2$ on GBP-R86N. Oxidized HEK293-GlycoDelete-produced GBP-R86N, a mixture of non-glycosylated (black, MW = 13469 Da), LacNAc (yellow, MW = 13834 Da) and oxidized LacNAc-Sia species (red, MW = 14063 Da), was ligated with biotin-PEG$_3$-ONH$_2$ in the presence of PPD both at 4°C (left) and 20°C (right). The conjugated species signal appears at the expected mass (green, MW = 14479 Da).

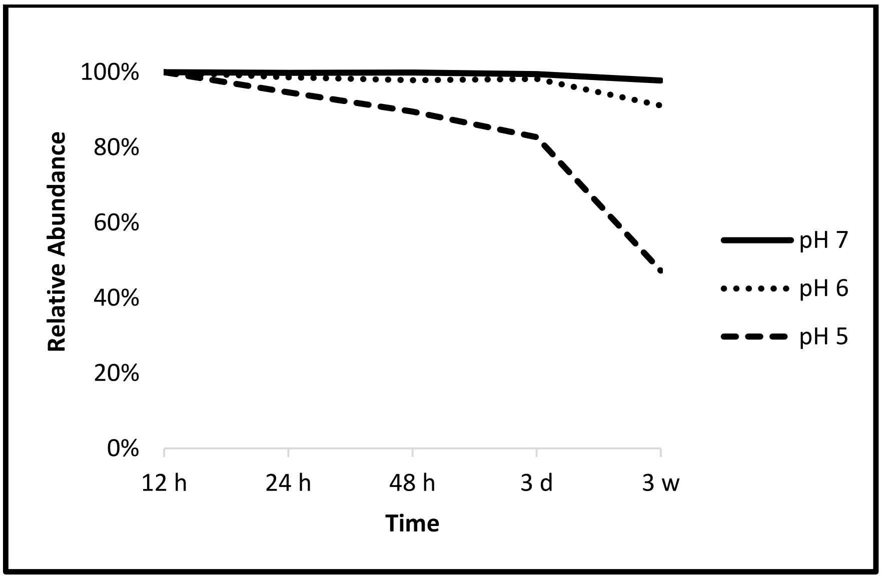


**Supplementary Figure 6.** Oxime stability at varying pH at 37°C. GBP-LacNAc-Sia-(oxime)-label was incubated at 37°C in buffers with varying pH (0.1 M Ac (pH 5), 0.1 M NaPi (pH 6) and 0.1 M NaPi (pH 7). Oxime hydrolysis was monitored at different time points.

### In Depth ESI-MS Analysis of a 5 kDa PEG Conjugation

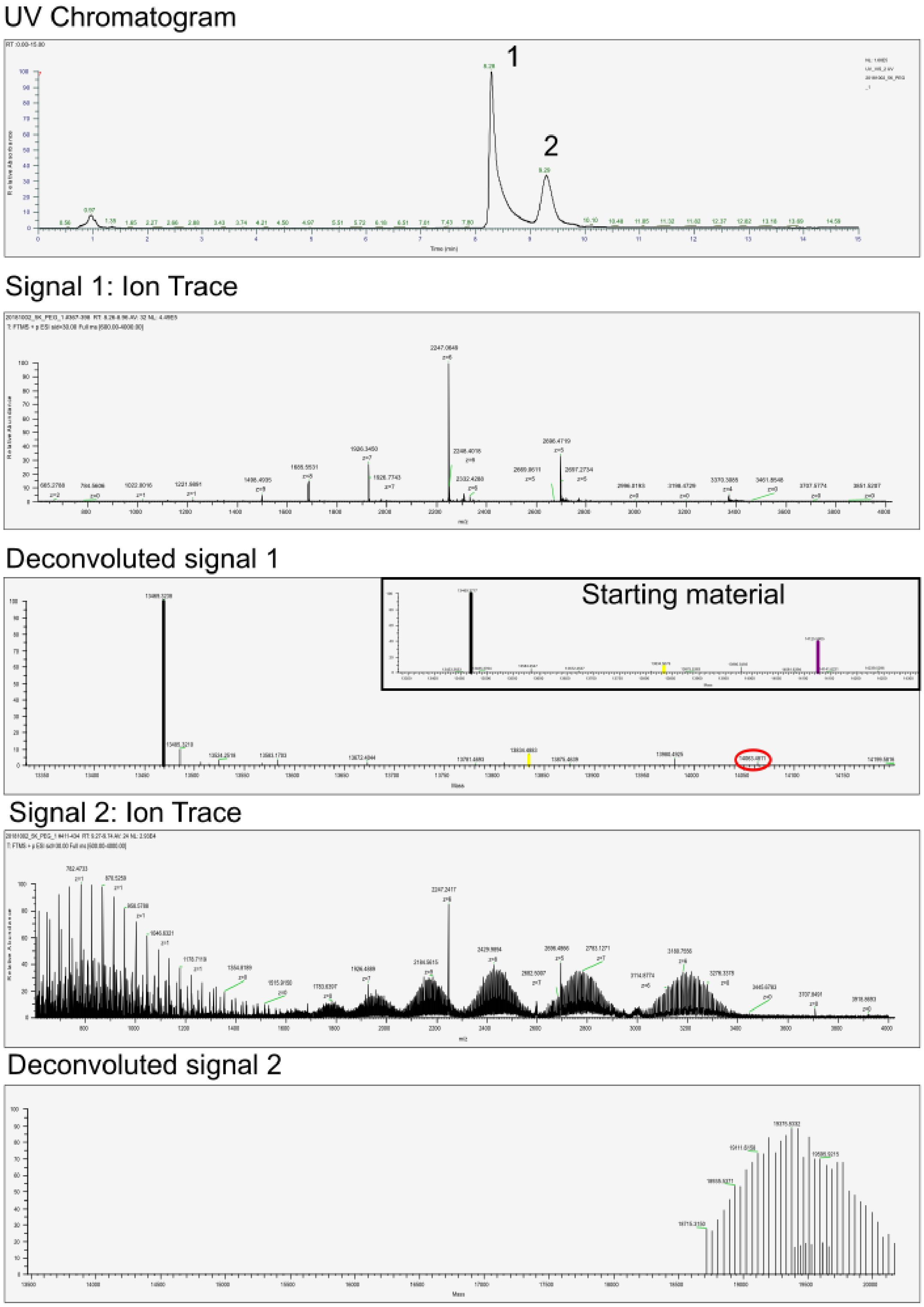

**Supplementary Figure 7**. In-depth MS analysis of the oxime-based 5-kDa PEG conjugation on NaIO4-oxidized GBP-R86N. The UV chromatogram (280 nm) shows two peaks eluting (signal 1 and 2). The MS spectrum of the first peak shows a typical pattern of a multi-charged protein. Upon deconvolution, the major mass that was found was that of non-glycosylated GBP-R86N (black, MW = 13469 Da). Next to that, only a small amount of oxidized LacNAc-Sia species was detected (red, MW = 14063 Da). The MS spectrum of the second peak shows a series of evenly spaced (44 Da) single charged masses distributed in a bell-shape. The 44 Da corresponds to the mass of a PEG monomer unit and the bell-shaped distribution is typical for the polydisperse nature of a polymer. These signals stem from leftover, free PEG polymer. Next to the free polymer, peaks corresponding to the conjugated protein can be recognised as densely packed, bell-shaped groups of signals. The deconvolution of these signals results in a evenly spaced (+44 Da) bell-shaped group of masses centred around 19 kD. This corresponds with the expected GBP-R86N-LacNAc-Sia-$PEG_n$ polymer conjugate (expected conjugate MW = 14062 Da + 5000 Da =19062 Da).

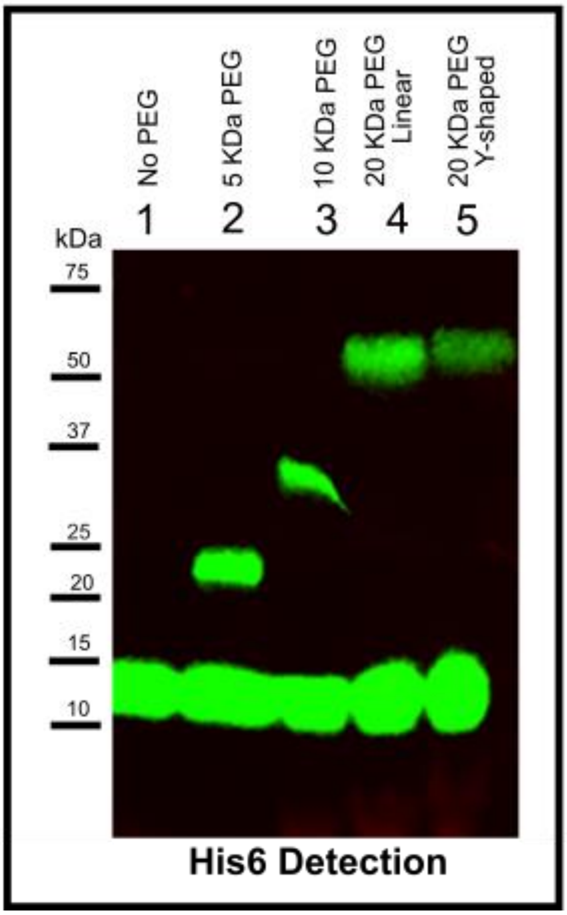


**Supplementary Figure 8.** Western Blot analysis of oxime-based PEG polymer conjugation on NaIO4-oxidized GBP-R86N. Oxidized HEK293-GlycoDelete-produced GBP-R86N, a mixture of non-glycosylated and LacNAc-Sia species, was conjugated with different PEG polymers (linear 5-kDa, 10-kDa and 20-kDa aminooxy-PEG polymer, and Y-shaped 20-kDa aminooxy PEG polymer).

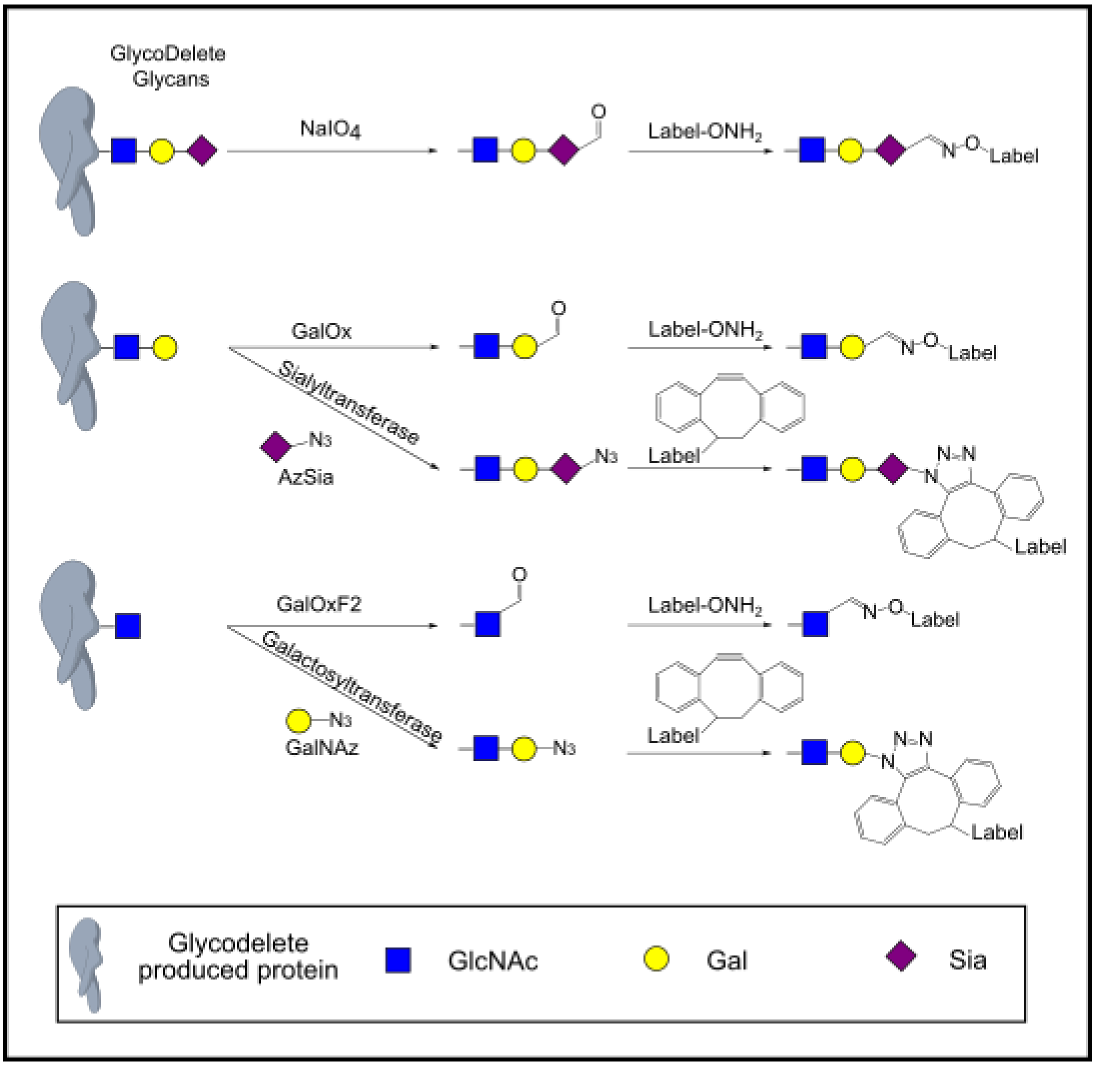


**Supplementary Figure 9.** Schematic overview of the different GlyConnect strategies explored in this work.
